# Fentanyl-induced antinociception, reward, reinforcement, and withdrawal in *Hnrnph1* mutant mice

**DOI:** 10.1101/2020.06.06.137158

**Authors:** Camron D. Bryant, Aidan F. Healy, Qiu T. Ruan, Michal A. Coehlo, Elijah Lustig, Neema Yazdani, Kimberly P. Luttik, Tori Tran, Isaiah Swancy, Lindsey W. Brewin, Melanie M. Chen, Karen K. Szumlinski

**Affiliations:** Laboratory of Addiction Genetics, Department of Pharmacology and Experimental Therapeutics and Psychiatry, Boston University School of Medicine; Department of Psychological and Brain Sciences, University of California, Santa Barbara; T32 Biomolecular Pharmacology Ph.D. Program, Boston University School of Medicine; Transformative Training Program in Addiction Science, Boston University; Undergraduate Research Opportunity Program (UROP), Boston University; Department of Molecular, Developmental and Cellular Biology and the Neuroscience Research Institute, University of California, Santa Barbara

**Keywords:** analgesia, addictive, pain, opiate, psychostimulant

## Abstract

Opioid Use Disorder (OUD) and opioid-related deaths remain a major public health concern in the United States. Both environmental and genetic factors influence risk for OUD. We previously identified *Hnrnph1* as a quantitative trait gene underlying the stimulant, rewarding, and reinforcing properties of methamphetamine. Prior work demonstrates that hnRNP H1, the RNA-binding protein encoded by *Hnrnph1,* post-transcriptionally regulates *Oprm1* (mu opioid receptor gene) – the primary molecular target for the therapeutic and addictive properties of opioids. Because genetic variants can exert pleiotropic effects on behaviors induced by multiple drugs of abuse, in the current study, we tested the hypothesis that *Hnrnph1* mutants would show reduced behavioral sensitivity to the mu opioid receptor agonist fentanyl. *Hnrnph1* mutants showed reduced sensitivity to fentanyl-induced locomotor activity, along with a female-specific reduction in, and a male-specific induction of, locomotor sensitization following three, daily injections (0.2 mg/kg, i.p.). *Hnrnph1* mutants also required a higher dose of fentanyl to exhibit opioid reward as measured via conditioned place preference. Male *Hnrnph1* mutants showed reduced fentanyl reinforcement. *Hnrnph1* mutants also showed reduced sucrose motivation, suggesting a reward deficit. No genotypic differences were observed in baseline thermal nociception, fentanyl-induced antinociception, physical or negative affective signs of opioid dependence, or in sensorimotor gating. In the context of our prior work, these findings suggest that *Hnrnph1* dysfunction exerts a selective role in reducing the addiction liability to drugs of abuse (opioids and psychostimulants), which could provide new biological pathways to improve their therapeutic profiles.

## INTRODUCTION

The United States is in the midst of a major opioid addiction epidemic, with over 2 million people estimated to be suffering from Opioid Use Disorder (**OUD**) and over 67,000 deaths in 2018 (https://www.cdc.gov). A major source of the current opioid epidemic was the over-prescribing of opioid analgesics that began in the late 1990s, based on the premise that pain is the fifth vital sign and that opioids are a safe and effective option with low risk for misuse if properly prescribed to pain patients. This philosophy led to a loosening of governmental regulations and opioid over-dispensing (Ostling *et al.* 2018). The recognition of a prescription opioid epidemic resulted in the advent of abuse-deterrent formulations of prescription opioids, such as Oxycontin^®^ (containing the mu opioid receptor agonist, oxycodone), which decreased the street value of heroin and made it more marketable (Cicero *et al.* 2012; Coplan *et al.* 2013). Recent laws limit the amount and frequency of opioid that can be dispensed within a prescription (e.g., Prescription Drug Monitoring Programs) (Fink *et al.* 2018) and have been associated with a massive shift to illicit heroin use over the past decade.

More recently, synthetic opioids, namely derivatives of the mu opioid receptor agonist fentanyl, have entered the market as they are synthesized more easily and affordably than heroin and can be more readily transported illicitly across borders, due to their increased potency. Thus, fentanyl-derivative drugs are frequently a major component of illicitly sold heroin. Fentanyl-derivative drugs can exhibit potencies ranging from 50 to 10,000 times greater than heroin or morphine, which makes a substantial contribution to the number of opioid-related deaths in the U.S. (Han *et al.* 2019).

While the increased availability of illicit prescription and non-prescription opioids has historically fueled the current opioid epidemic, other environmental risk factors (e.g., early life stress) (Webster 2017) and genetic factors (Jensen 2016) also contribute to risk for OUD. Twin and family studies estimate the heritability of OUD at greater than 50% (Goldman *et al.* 2005; Ho *et al.* 2010). However, the genetic basis of OUD remains largely unknown, with only a handful of genome-wide association studies (**GWAS**) reporting genome-wide significant loci associated with addiction, including *KCNC1* and *KCNG2* (voltage-dependent potassium channel and channel modifier, respectively), *CNIH3* (AMPA receptor axillary protein 3), and most recently *RGMA* (RGM domain family, member 3) (Jensen 2016; Nelson *et al.* 2016; Berrettini 2017; Cheng *et al.* 2018). Interestingly, a genome-wide association study identified a noncoding single nucleotide polymorphism (SNP) upstream of *OPRM1* (opioid receptor, mu 1) that was associated with the therapeutic dose of methadone for treating OUD (Smith *et al.* 2017). Larger sample sizes, including an increase in the number of opioid-exposed unaffected controls, will yield several new genome-wide discoveries in the coming years (Berrettini, 2017).

RNA binding proteins (**RBPs**) are a diverse class of molecules that bind to RNA and regulate all aspects of RNA metabolism, including post-transcriptional processing, transport, cellular localization, and local translation. RBPs, including heterogeneous nuclear ribonuclear proteins (**hnRNPs**), can translocate to the cytoplasm following exposure to various extracellular stimuli (e.g., stressors and neuronal activation) and regulate local translation underlying activity-dependent synaptic plasticity (Zhang *et al.* 2012). Acute and repeated exposure to drugs of abuse induces activity-dependent synaptic plasticity within the limbic system, including the mesocorticolimbic circuit that underlies behavioral manifestation of the addictions (Luscher & Malenka 2011). Because RBPs are positioned to locally and rapidly regulate synaptic protein translation following drug-induced modulation of cellular activity and to regulate long-term drug-induced changes in nuclear gene transcription, there is a growing appreciation that RBPs likely play a pivotal role in synaptic plasticity underlying addiction (Bryant & Yazdani 2016).

We used an unbiased, forward genetic and fine mapping approach to positionally clone and validate the RBP heterogeneous nuclear ribonuclear protein H1 *(**Hnrnph1**)* as a quantitative trait gene underlying the locomotor stimulant response to methamphetamine (Yazdani *et al.* 2015). Differential exon usage analysis identified a set of four quantitative trait variants within the 5’ UTR of *Hnrnph1* that decrease 5’ UTR usage, decrease hnRNP H protein expression, and functionally decrease luciferase reporter expression (Ruan *et al.* 2020b). *Hnrnph1* mutants harboring a frameshift mutation in the first coding exon show decreased methamphetamine-induced reward, reinforcement, and dopamine release (Ruan *et al.* 2020a). Furthermore, D1 dopamine receptor activation caused an increase in nuclear hnRNP H immunofluorescence in primary rat cortical neurons that was blocked with a D1 dopamine receptor antagonist (Ruan *et al.* 2018), suggesting post-synaptic modulation of hnRNP H in response to D1 receptor signaling.

With regard to opioids, *HNRNPH1* contributes to post-transcriptional processing of *Oprm1,* the gene encoding the mu opioid receptor, including 5’ UTR-mediated translational repression (Song *et al.* 2012) and splicing (Xu *et al.* 2014). The mu opioid receptor is the primary molecular target underlying the addictive and therapeutic properties of opioid drugs (Matthes *et al.* 1996). In support of a potential role for *Hnrnph1* in OUD, an intronic variant in *OPRM1* that affects hnRNP H1 binding was associated with the severity of heroin dependence and alternative splicing of *OPRM1* in humans (Xu *et al.* 2014).

Genetic loci and genetically engineered mutations can exert pleiotropic behavioral effects on multiple classes of drugs of abuse (Tammimäki & Männistö 2011), including psychostimulants and opioids (Bryant *et al.* 2009b, 2009a, 2012; Goldberg *et al.* 2017). Thus, in the present study, we examined OUD-related traits in *Hnrnph1* mutant mice that display deficits in methamphetamine-induced locomotor activity, reward, reinforcement, and dopamine release (Ruan *et al.* 2020a). We employed fentanyl as our mu opioid receptor agonist of choice, given its high degree of selectivity for the mu opioid receptor, its rapid onset of action that can facilitate drug-related learning, and the global prevalence of abuse and deaths associated from its illicit use (Han *et al.* 2019). Because both psychostimulant- and opioid-induced locomotor activity depend on dopamine release in the striatum (Wise & Bozarth 1987; Di Chiara & Imperato 1988; Di Chiara & North 1992), we hypothesized that *Hnrnph1* mutants would display reduced sensitivity to fentanyl-induced locomotor activity and perhaps other behaviors related to drug reward and reinforcement.

Herein, we examined a large battery of behavioral traits that model various aspects of OUD, including acute sensitivity and sensitization to fentanyl-induced locomotor stimulation, conditioned and state-dependent reward as measured via conditioned place-preference, reinforcement as measured via oral self-administration under operant-conditioning procedures, acute antinociception and tolerance, as well as affective and physiological signs of drug dependence during fentanyl withdrawal. Importantly, we included sex as a biological variable to examine potential Genotype × Sex interactions in mediating the behavioral effects of *Hnrnph1* deletion on OUD-relevant behaviors. These results identify select, sometimes sex-dependent, changes in fentanyl-induced behavioral phenotypes in *Hnrnph1* mutant mice.

## MATERIALS AND METHODS

### Mice

*Hnrnph1* mutant mice *(**Hn1+/−**)* were generated using TALENs targeting the first coding exon of *Hnrnph1* (exon 4, UCSC Genome Browser; https://genome.ucsc.edu/), resulting in a frameshift mutation and a premature stop codon. *Hn1*+/− mice show reduced transcription of the wild-type (**WT**) *Hnmph1* transcript, an upregulation of the mutant and total transcript levels (Yazdani *et al.* 2015), and a two-fold increase in hnRNP H striatal synatosomal protein (Ruan *et al.* 2020a). Therefore, *Hn1*+/− refers to the TALENs-induced indel in exon 4 that leads to a premature stop codon and not to gene haploinsufficiency.

Experimental mice were generated by mating *Hn1*+/− males with C57BL/6J females purchased from The Jackson Laboratory (Bar Harbor, ME, USA for studies conducted at Boston University; Sacramento, CA, USA for studies conducted at the University of California Santa Barbara; UCSB). Offspring were genotyped and, unless otherwise indicated, female and male littermates from a minimum of five different litters, ranging from 56-100 days of age, were employed in the studies. Mice were housed in same-sex groups of 3-5 in standard mouse cages, housed within ventilated racks under standard housing conditions. Mice involved in the fentanyl self-administration experiments were housed under a reverse light cycle (lights off: 1000 h), otherwise, all other mice were housed under a regular 12 h/12h light/dark cycle (lights on: 0700 h at UCSB; lights on: 06:30 at Boston University). All experiments were conducted in compliance with the National Institutes of Health Guide for Care and Use of Laboratory Animals (NIH Publication No. 80-23, revised 2014) and approved by the IACUCs of UCSB and Boston University.

We used a previously published power analysis (Ruan *et al.* 2020a) based on our original finding of decreased MA-induced locomotor activity in *Hn1*+/− mice (Yazdani *et al.* 2015) to guide selection of our sample size. Briefly, with an effect size of Cohen’s d = 0.9, we used G*Power 3 (Faul *et al.* 2007) and determined that a sample size of n = 16 per genotype is required to achieve 80% power (p < 0.05).

### Drugs

Fentanyl citrate was purchased from Sigma-Aldrich (St. Louis, MO USA) and was dissolved in warm physiological saline (0.9% NaCl) for intraperitoneal (i.p.) injection or in tap water for oral consumption. Buspirone hydrochloride (Sigma-Aldrich) and naltrexone (Tocris Bioscience/Bio-Techne (Minneapolis, MN, USA) were dissolved in sterile saline for i.p. injection.

### Fentanyl locomotor activity and sensitization

On Days 1 and 2 of locomotor testing, mice received a saline injection (10 ml/kg) and locomotor activity was recorded for 30 min. On Days 3-5, mice received a fentanyl injection (0.2 mg/kg) and locomotor activity was recorded for 30 min each day. The dose of fentanyl was chosen based on previous studies indicating a robust increase in locomotor activity in C57BL/6J mice (Bryant *et al.* 2009a, 2009c, 2012; Goldberg *et al.* 2017). Mice were recorded with infrared security cameras (Swann Communications, U.S.A., Inc., Santa Fe Springs, CA USA) mounted above the Plexiglas chambers [40 cm long × 20 cm wide by 45 cm high (Yazdani *et al.* 2015)]. Data analyses were performed in R (https://www.r-project.org/).

### Fentanyl conditioned place preference

Experimentally naïve mice were tested for fentanyl-conditioned place-preference (**CPP**). The same locomotor apparatus was partitioned into two equal-sized compartments via a black, ion transparent, plastic divider with a mouse entryway (5 cm × 6.25 cm) that was flipped upside down during training and was used to confine mice to one side. Behavior was recorded using digital video-tracking (Anymaze, Stoelting Co., Wood Dale, IL USA). A 9-day CPP protocol was employed that included 30-min conditioning and test sessions (Yazdani *et al.* 2015). On Day 1, initial preference was assessed whereby mice received an injection of saline (10 ml/kg, i.p.), were placed on the left side, and were provided open-access to both sides. On conditioning Days 2 and 4, all mice received saline and were confined to the left side that contained smooth plastic floor inserts (Plaskolite). On conditioning Days 3 and 5, mice received either saline, 0.05 mg/kg, or 0.2 mg/kg fentanyl and were confined to the right side that contained a textured plastic floor insert. On Days 6 and 7, mice were left undisturbed in their home cages. On Day 8, mice received an injection of saline, were placed into the left side, and were provided open-access to both sides for 30 min. The difference in time spent on the drug-paired side between Day 8 and Day 1 was calculated to index the magnitude of the conditioned response. On Day 9, mice received a priming injection of either saline or their fentanyl-conditioning dose, were placed into the left side, and provided openaccess to both sides. The difference in time spent on drug-paired side between Day 9 and Day 1 was calculated and indexed the magnitude of conditioning in a fentanyl-dependent state (Kirkpatrick & Bryant 2015).

### Baseline nociception and fentanyl antinociception

The mice used to assess the acute antinociceptive effects of fentanyl served previously as the saline control group in the CPP experiment. Thus, these mice had a history of five saline injections, but were completely opioid-naïve. For testing of baseline nociception and fentanyl antinociception, the hot plate temperature was set to 52.5°C. Mice were habituated to the testing room for at least 1 h. Mice were then placed in a Plexiglas cylinder (15 cm diameter; 33 cm in height) on the hot plate (IITC Life Science Inc.) and two separate baseline latencies to lick the hind paw were recorded, separated by 30 min. Thirty min post-baseline assessment, mice were injected with cumulative doses of fentanyl (0.1, 0.1, 0.2, and 0.4 mg/kg) and tested for post-injection hind pawlick latencies every 10 min. Within each trial, a cut-off latency of 60 s was employed to avoid tissue damage. Percent maximum possible effect (% MPE) was calculated using the following formula: %MPE = (post-injection latency minus pre-injection latency) / (60 – pre-injection latency) * 100.

In an attempt to induce antinociceptive tolerance, we treated a separate, experimentally naïve cohort of mice with repeated injections twice daily for five days [0.8 mg/kg fentanyl (i.p.) or saline (i.p.); 0900 h and 1700 h] for 5 consecutive days and then tested with a challenge, submaximal antinociceptive dose of 0.4 mg/kg fentanyl (i.p.) on the sixth day, beginning at 0900 h (**see Supplementary Information for additional details**).

### Fentanyl operant conditioning

Male mice were first trained to lever press for delivery of a 10% sucrose solution under a fixed ratio 1 (**FR1**) schedule of reinforcement with a 20 s timeout in standard mouse operant chambers (Med Associates, St. Albans, VT USA). Male mice were employed exclusively in this initial operant-conditioning study due to limited availability of female mice at the time the experiment was performed. As in recent studies (Szumlinski *et al.* 2019), each right lever-press resulted in delivery of 20 μl of the sucrose solution and a 20 s presentation of a tone/light stimulus-complex. Left lever presses resulted in no programed consequences. Sucrose training proceeded for 12 days, by which point, mice of both genotypes had exceeded the minimum requirements for successful acquisition of the operant response (a minimum of 10 active lever-presses in 60 min + greater than 70% responding on the active lever for 3 consecutive days). Then, the sucrose solution was substituted for an unadulterated 3 mg/L fentanyl solution – a concentration demonstrated recently to be readily consumed by mice under free-access conditions in the home-cage (Szumlinski *et al.* 2019) – and mice underwent daily, 60 min testing sessions for an additional 10 days. In the initial fentanyl self-administration study, the concentration of the fentanyl reinforcer was then progressively *increased* across days (10, 30, 100, 300 and 1000 mg/L), with each concentration presented until responding stabilized (less than 15% variability across three consecutive presentations) or for a maximum of 10 days of self-administration.

Because the data from this initial fentanyl experiment suggested that the doses tested were located on the descending limb of the dose-response function (see Results), we trained a separate cohort of female and male mice to nose-poke for 3 mg/L fentanyl during once daily (60-min) sessions, in the absence of any prior sucrose-training. In this second study, mice of both genotypes met the acquisition criterion for self-administration training within 10 days, at which time the fentanyl concentration available was progressively *decreased* across days (1,0.3, 0.1 and 0.03 mg/L), with each concentration presented until responding stabilized or for a maximum of 10 days of self-administration.

### Fentanyl withdrawal-induced negative affect and physical dependence

To examine the effects of *Hn1*+/− on fentanyl withdrawal-induced negative affect and physical dependence, female and male *Hn1*+/− and wild-type (**WT**) littermates were injected twice daily (0900 h and 1700 h) with 0.8 mg/kg fentanyl (i.p.) or saline (i.p.) for 5 days. The next morning, mice were subjected to a 1-day behavioral test battery to assess fentanyl withdrawal-induced sensorimotor-gating deficits and negative affect. The 1-day test battery was very similar to a battery that we employed in a recent opioid study (Szumlinski *et al.* 2019), as well as prior studies of alcohol withdrawal-induced negative affect (Lee *et al.* 2016, 2017a, 2018). The battery consisted of testing for acoustic startle and prepulse inhibition of acoustic startle, followed, in order, by testing in the light-dark shuttle box test, novel object encounter, marble-burying and the Porsolt forced swim tests. Details of the specific procedures are provided in the **Supplementary Information**. The day following the behavioral test battery, fentanyl-experienced mice were then injected in the morning (~0800 h) with 0.8 mg/kg fentanyl. Eight hours later, mice were injected with 10 mg/kg naltrexone and tested for physiological signs of withdrawal as described previously (Jimenez *et al.* 2017; Szumlinski *et al.* 2019) and detailed in the **Supplementary Information**. To accommodate all the mice, behavioral testing was conducted in cohorts of 8 mice, with ~24 mice tested per day.

### Statistical analysis

Locomotor activity was analyzed in 5 min bins in R (https://www.r-project.org/) using a mixed-design Genotype × Sex × Day ANOVA, with Day as the repeated measure. In the cases where Genotype × Sex interactions were detected, the interaction was deconstructed along the Sex factor for follow-up analyses using t-tests, based on main effects and interaction terms. Locomotor habituation to saline injections was quantified as change in locomotor activity on Day 1 versus Day 2 (Day 1 – Day 2). Taking into account genotypic differences in baseline locomotion, we also quantified the acute locomotor response to fentanyl as the change in locomotor activity on Day 3 (first fentanyl injection) versus Day 2 (second saline injection). We also quantified locomotor sensitization to fentanyl as the change in locomotor activity from Day 3 to Day 5 (i.e., the first to third fentanyl injection).

For the conditioning experiments, data analyses were performed in R. Three-way ANOVAs (Genotype, Sex, Dose) were employed. For nociception, fentanyl antinociception, and tolerance, data analyses were also performed in R. We first analyzed acute fentanyl antinociception via a mixed model ANOVA with Genotype and Sex as factors and Cumulative Dose as the repeated measure.

For operant conditioning, analyses were performed using SPSS v.21 statistical software (IBM, 2012). The average number of active and inactive responses, the average response allocation on the fentanyl-reinforced operandum (i.e., percentage of total responses directed at the lever or hole that resulted in reinforcer delivery) and the average intake of sucrose or fentanyl during the initial training phase and during the last 3 days of the training phase were analyzed using t-tests or Genotype × Sex ANOVAs, as appropriate. The data from the dose-response phase of testing were analyzed using a Genotype × Dose ANOVA (experiment 1) or Genotype × Sex × Dose ANOVA (experiment 2), with Dose as a repeated measure.

For fentanyl withdrawal and physical dependence, unless otherwise indicated, the data were analyzed using a Genotype × Sex univariate ANOVA using SPSS v.23 software.

## RESULTS

### Hn1+/− decreases acute fentanyl locomotor activity, decreases fentanyl locomotor sensitization in females, and increases fentanyl locomotor sensitization in males

Using a within-subjects design, we first examined the effects of *Hnrnph1* deletion on the locomotor response to saline (i.p.; Days 1, 2) and fentanyl (0.2 mg/kg, i.p.; Days 3, 4, and 5). There was a significant Genotype × Sex × Day interaction (F_4,232_ = 3.92, p = 0.004) (**Fig.1A**) and thus, the interaction was deconstructed along the Sex factor to examine for sex differences in the effects of *Hn1*+/− on behavior.

**Figure 1.**
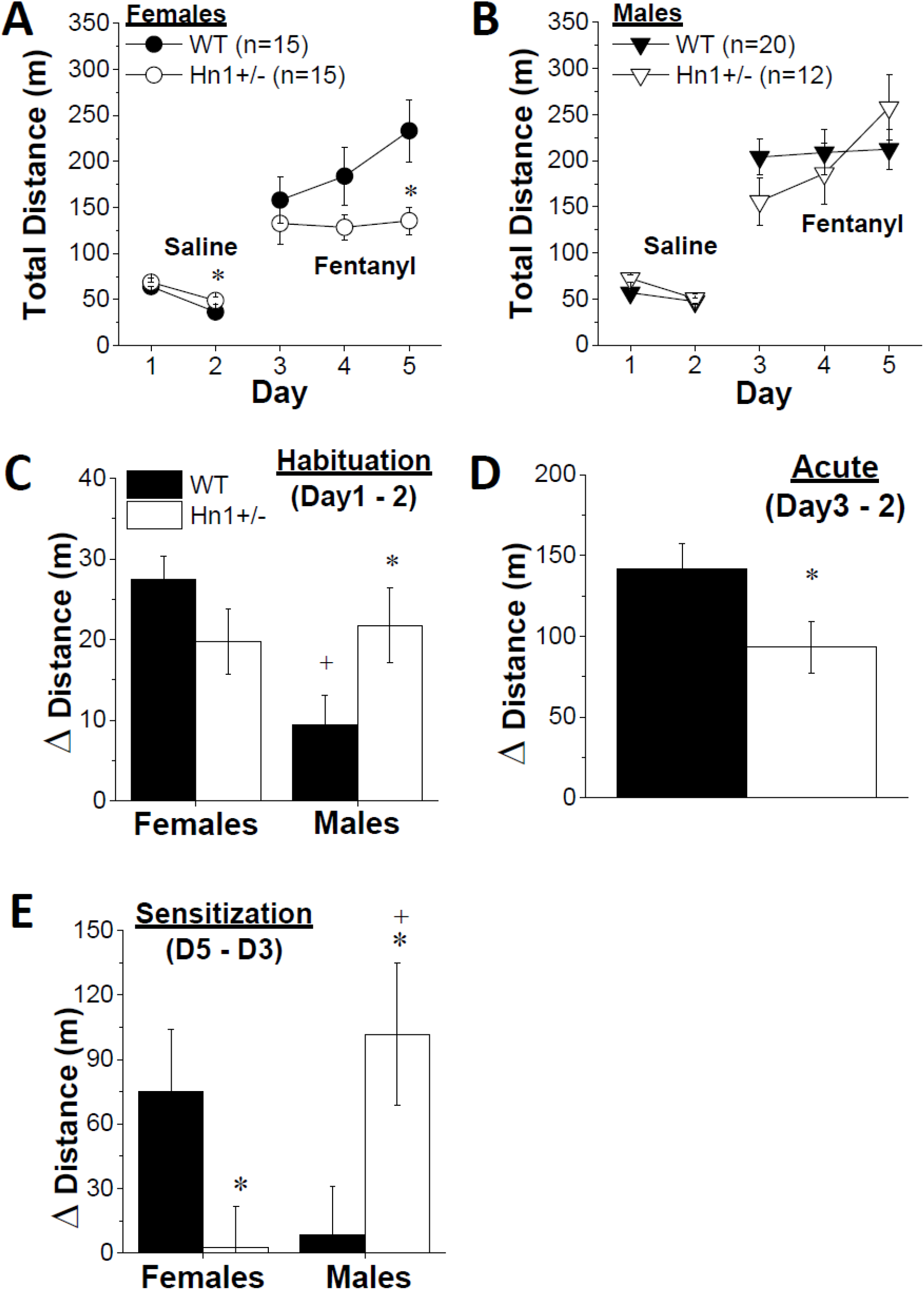
Sex-dependent modulation of fentanyl-induced locomotor activity and sensitization in *H1*+/− mice. **(A):** In response to saline (i.p.), *Hn1*+/− females showed greater saline-induced locomotor activity than their WT controls following the 2^nd^ saline injection (Day 2; *p = 0.028). In response to fentanyl, *Hn1*+/− females showed reduced fentanyl-induced locomotor activity following the 3^rd^ injection of 0.2 mg/kg fentanyl (i.p.; Day 5; *p = 0.014). **(B):** In contrast no genotypic differences in saline- or fentanyl-induced locomotion were observed in males. **(C):** In WT mice, the locomotor response to repeated saline (Day 1 – Day 2) habituated to a lesser extent in males versus females (**+**p = 0.0008), while no sex difference was apparent in *Hn1*+/− mice. Further, saline-induced locomotion showed greater habituation in *Hn1* +/− males relative to WT males (*p = 0.047). No genotypic difference in habituation was observed for females. **(D):** When taking into account genotypic differences in baseline locomotor activity (i.e. response to the 2^nd^ saline injection), *Hn1*+/− mice showed less robust acute fentanyl-induced locomotor activity (0.2 mg/kg, i.p.) compared to WT mice (**\***p = 0.039). **(E):** A marked Genotype × Sex interaction was apparent with respect to the change in fentanyl-induced locomotion across the 3 injections, with *Hn1*+/− females showing less (**\***p = 0.04) and *Hn1*+/− males showing more (**\***p = 0.02) sensitization compared to their female and male WT counterparts, respectively. Data represent the mean ± S.E.M. Ns are included in the figure legends. *p < 0.05 vs. WT, **+**p < 0.05 vs. other sex.

A significant Genotype × Day interaction was detected in females (F_4,112_=4.15, p=0.004) that reflected increased locomotor activity in *Hn1*+/− females following the second saline injection on Day 2 (t_28_ = 2.32, **\***p = 0.028) and decreased fentanyl-induced locomotor activity in *Hn1*+/− females on Day 5 (the 3^rd^ fentanyl injection; t_28_ = 2.63, *p = 0.014; **Fig. 1A**). Although fentanyl administration resulted in increased locomotor activity in males (Day effect: F_4,120_=48.62, p<0.0001), no genotypic differences were detected (Genotype effect and interaction, p’s>0.1; **Fig.1B**).

In examining habituation to saline-induced locomotor activity (activity on Day 1 – Day 2), we detected another Genotype × Sex interaction (F_1,58_ = 6.69, p = 0.012) that reflected less habituation in WT males compared to WT females (**Fig.1C:** t_33_ = 3.66; **+**p = 0.0008) and more habituation in *Hn1+/−* males versus WT males (**Fig.1C:** t_30_ = 2.076, *p=0.047), with no genotypic difference observed in females (t_33_ = 1.56, p = 0.13). Sex did not influence the locomotor response to acute fentanyl in either genotype (Day 3 – Day 2; Sex effect: F_1,58_ = 1.65, p = 0.20; interaction: F_1,58_ < 1). However, a significant Genotype effect was detected for the Day 3 – Day 2 phenotype (F_1,58_ = 4.45, **\***p = 0.039) that reflected lower fentanyl-induced locomotor activity in *Hn1*+/− versus WT mice (**Fig.1D**).

In examining the extent to which the second and third fentanyl injections on Days 4 and 5 induced locomotor sensitization relative to the first fentanyl injection on Day 3 (Day 5 – Day 3), there was a robust Genotype × Sex interaction (F_1,58_ = 10.15, p = 0.002) that reflected less sensitization in *Hn1+/−* females and more sensitization in *Hn1*+/− males versus their respective WT counterparts (females: t_30_ = 2.11, p = 0.04; males: t_30_ = 2.40, p = 0.02; **Fig.1E**). Thus, while the effect of *Hn1*+/− on the acute locomotor response to fentanyl is sex-independent, the regulation of fentanyl-induced locomotor sensitization varies markedly as a function of Sex.

### Reduced fentanyl-CPP in Hn1+/− mice

In examining drug-free expression of fentanyl reward via CPP (Day 8 – Day 1), there was a main Dose effect (F_2,136_ = 3.32, p = 0.039), but no Genotype effect (F_1,136_ < 1) or Sex effect (F_1,136_ = 2.48, p=0.12). Importantly, there was a significant Genotype × Dose interaction (F_2,136_ = 3.46, p=0.034) that was partially explained by WT mice showing a significant fentanyl-CPP at 0.05 mg/kg fentanyl compared to their WT saline (0 mg/kg) counterparts [t_46_ = 2.07; **#**p = 0.044] and by *Hn1*+/− mice showing a significant fentanyl-CPP at 0.2 mg/kg fentanyl relative to their *Hn1*+/− saline (0 mg/kg) counterparts [t_45_ = 2.67; **#**p = 0.011; **Fig.2A**). The Genotype × Dose interaction was also partially explained by *Hn1+/−* mice showing more fentanyl-CPP compared to WT mice at 0.2 mg/kg (t_46_ = 3.07; **\***p = 0.0036).

**Figure 2.**
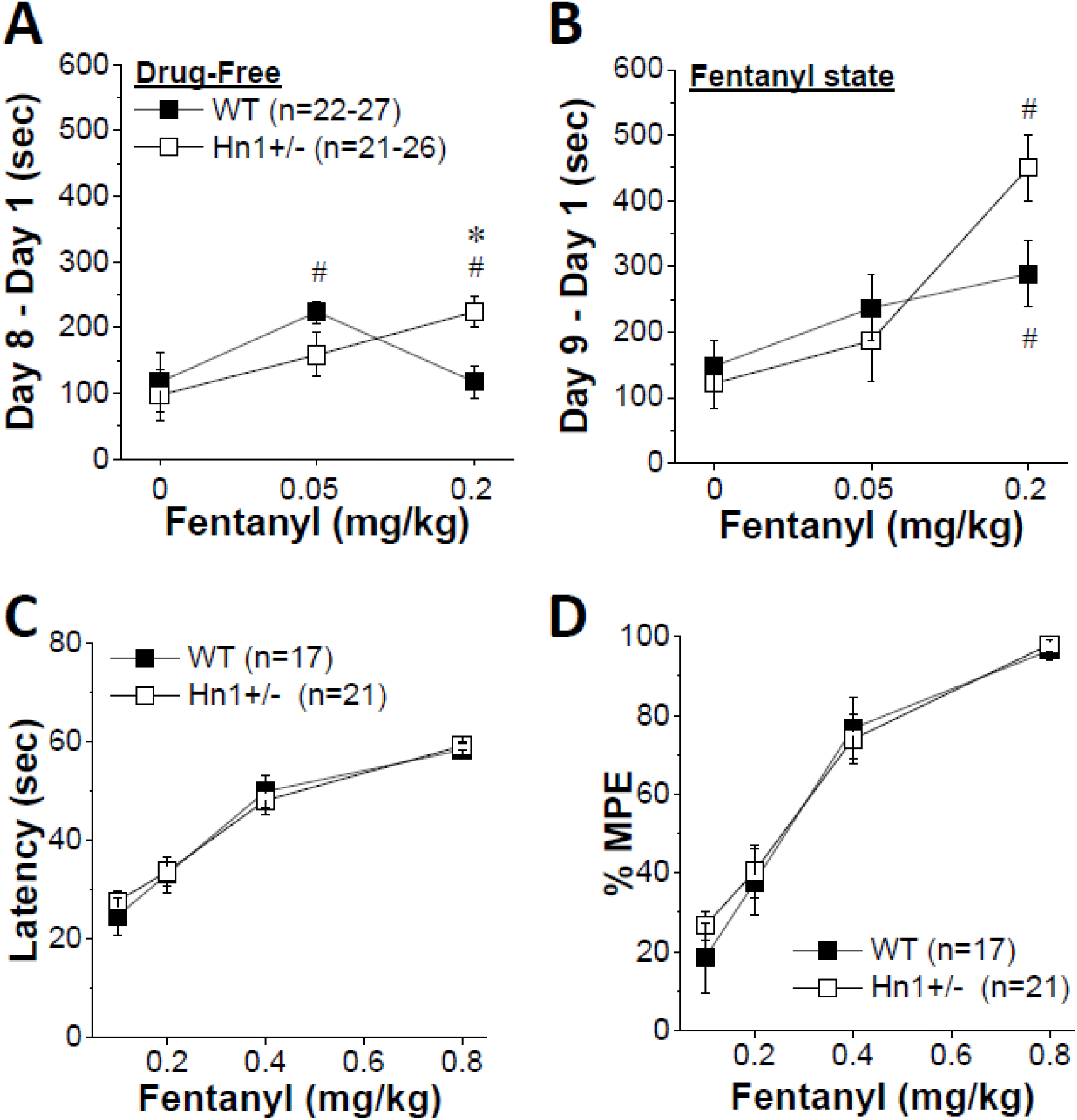
Modulation of fentanyl reward but not antinociception in *H1*+/− mice. **(A):** When assessed in a drug-free state, WT mice conditioned with 0.5 mg/kg fentanyl exhibited a significantly greater conditioned place preference (CPP) than mice conditioned with saline (versus 0 mg/kg; **#**p = 0.044), while significant CPP was evident in *Hn1*+/− mice only with the 2.0 mg/kg dose (versus 0 mg/kg; **#**p = 0.011). **(B):** When tested under the influence of their conditioning dose, both WT and *Hn1*+/− mice exhibited a significant CPP at the 2.0 mg/kg fentanyl dose (versus 0 mg/kg; **#**p = 0.033, < 0.0001, respectively). **(C):** No genotypic difference was apparent for the latency to lick the hind paw at baseline or in fentanyl antinociception following treatment with a cumulative fentanyl-dosing regimen. **(D):** Likewise, no genotypic difference was noted in the precent maximum possible effect (%MPE) of fentanyl antinociception. Data represent the mean ± S.E.M. of the number of mice indicated in the figure legends. **#**p < 0.05 versus 0 mg/kg fentanyl.

When mice were tested for state-dependent CPP in a fentanyl-primed state (i.e., under the influence of their respective training dose), there was a significant Dose effect (F_2,136_ =10.98, p < 0.0001) and Sex effect (F_1,136_ = 4.36, p = 0.039), but no Genotype effect (F_1,136_ <1) or interactions (all p’s > 0.075). As illustrated in **Fig.2B**, both WT mice (t_51_ = 2.19, **#**p = 0.033) and *Hn1*+/− mice (t_45_ = 5.33, **#**p < 0.0001) primed with the 0.2 mg/kg fentanyl dose exhibited a robust place-preference, relative to their 0 mg/kg counterparts primed with a saline injection.

In examining locomotor activity during fentanyl-CPP assessment (Days 1, 8, and 9) and during fentanyl-CPP training (Days 2, 3, 4, and 5), we found little evidence for an effect of *Hn1*+/− on drug behavior in this context (**Supplementary Figures 1 and 2**).

### No change in baseilne thermal nociception or fentanyl antinociception following acute or repeated fentanyl administration in Hn1+/− mice

In examining the first, second, and average baseline hot plate latencies, there was no significant effect of Genotype (F_1,34_ < 1 for all three measures), Sex (F_1,34_ = 2.05, 1.12, 1.90; p = 0.16, 0.30, 0.18), or interaction (F_1,34_ = 3.32, 0.083, 1.36; p = 0.077, 0.78, 0.25). Furthermore, there was no significant genotypic difference in acute fentanyl-induced antinociception following a cumulative dosing procedure (0-0.8 mg/kg, i.p.) and assessment of post-injection latencies (Genotype and Sex effects: F_1,34_ < 1; Dose effect: F_3,102_ = 98.19, p < 0.0001; all interactions involving Genotype: p’s > 0.54; **Fig.2C**) or in assessment of % MPE (Genotype and Sex effects: F_1,34_ < 1; Dose effect: F_3,102_ = 95.68, p < 0.0001; all interactions involving Genotype: p’s > 0.36; **Fig.2D**). These data fail to support an effect of *Hn1*+/− on acute fentanyl-induced antinociception.

In examining fentanyl-induced antinociception in *Hn1*+/− versus WT littermates following repeated, twice-daily injections of fentanyl (0.8 mg/kg, i.p.), there was no significant effect of *Hn1*+/− on baseline nociception (first, second, averaged) or on fentanyl-induced antinociception (0.4 mg/kg, i.p.), regardless of Chronic Treatment (**Supplementary Figure 3**).

### Decreased sucrose operant responding in Hn1+/− males

In examining operant-responding for reinforcement for 10% (w/v) sucrose in males, on average, *Hn1*+/− males showed significantly fewer active lever-presses than WT males during the 12-day training period (**Fig. 3A**) (t_22_ = 2.93, **\***p = 0.008). Although both genotypes met the acquisition criterion for sucrose self-administration by the end of the training period, active lever-responding was still significantly lower in *Hn1*+/− males versus WT males over the last 3 days of training (t_22_ = 2.99, p = 0.008; WT = 73.73 ± 15.50 vs. *Hn1*+/− = 25.57 ± 8.01). Similarly, *Hn1*+/− males also emitted fewer inactive lever-presses during the sucrose phase of the study, as evidenced both in terms of the average overall number of inactive lever-responses (**Fig. 3B**) (t_22_ = 2.56, **\***p = 0.02) and the average number of inactive lever-presses during the last 3 days of sucrose training (t_22_ = 2.63, p = 0.02; WT = 19.87 ± 6.48 vs. *Hn1*+/− = 4.55 ± 1.78). Importantly, there were no genotypic differences in the allocation of responding towards the sucrose-reinforced lever, indicating that *Hn1*+/− deletion did not significantly impair sucrose reinforcement, at least in males (**Fig. 3C**) (for average of the 12-day training period: t_22_ = 0.12, p = 0.90; for the last 3 days of sucrose training: t_22_ = 0.09, p = 0.93; WT = 73.18 ± 6.83% vs. *Hn1*+/− = 72.21 ± 8.42%). Finally, while the average sucrose intake was lower in *Hn1*+/− males versus WT males, this difference was not statistically significant (**Fig. 3D**) (for average of the 12-day training period: t_22_ = 1.89, p = 0.12; for the last 3 days of sucrose training: t_22_ = 1.57; p = 0.23; WT = 2.51 ± 0.62 g/kg vs. *Hn1*+/− = 1.87 ± 0.45 g/kg). These data provide new evidence that *Hnrnph1* gene products regulate behavioral output of male mice under operant-conditioning procedures but are less involved in regulating the appetitive and consummatory aspects of sucrose reinforcement.

**Figure 3.**
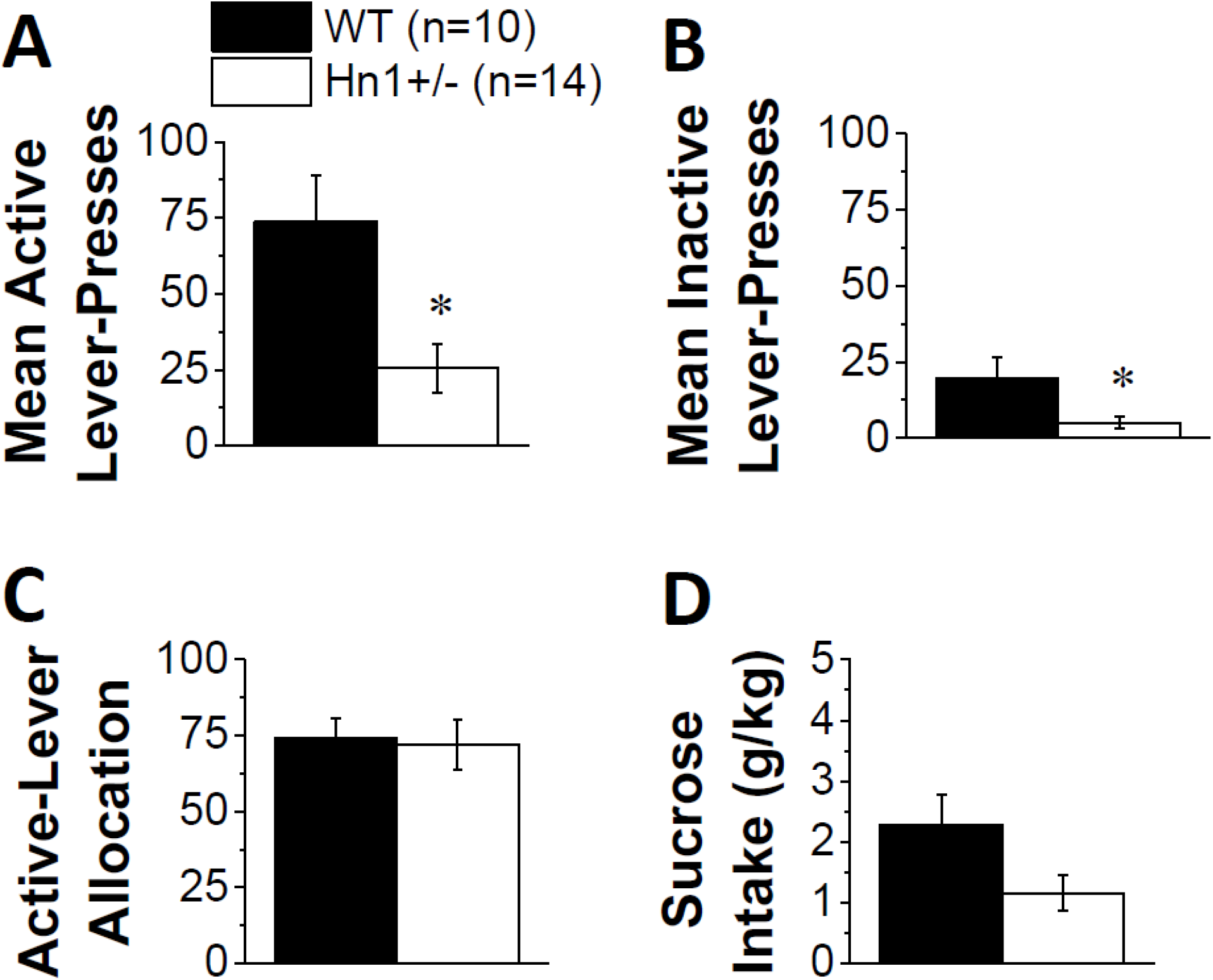
Blunting of operant-responding for sucrose reinforcement in *Hn1*+/− mice. When trained to leverpress for 10% sucrose, *Hn1*+/− males exhibited fewer active (**A: \***p = 0.008) and inactive (**B: \***p = 0.02) leverpresses, than WT controls. However, both genotypes directed a similar percentage of their responses toward the sucrose-reinforced lever, indicating that the mutation did not alter fentanyl-directed responding (**C**). **(D):** While sucrose intake was lower in *Hn1*+/− versus WT males, this genotypic difference was not statistically significant. Data represent the mean ± S.E.M. Ns are indicated in the figure legends. **\***p < 0.05 vs. WT

### Decrease in operant fentanyl intake in Hn1+/− males

As detailed in the Supplemental Results, when sucrose-trained mice were subsequently trained to lever-press for oral fentanyl reinforcement (3 mg/L), *Hn1+/−* males showed reduced indices of fentanyl reinforcement during the 10-day training phase of the study (**Supplementary Figure 4A-D**). However, no significant *Hn1+/−* effect was observed for the dose-response functions related to high-dose (3-1000 mg/L) fentanyl reinforcement (**Supplementary Figure 4E-H**). As also detailed in the Supplemental Results, when fentanyl reinforcement was assessed a second time in sucrose-naïve female and male mice, there were no significant genotypic differences in drug self-administration or intake during the training phase of the study (i.e., when 3 mg/L fentanyl served as the reinforcer), although males emitted more active nose-pokes and exhibited greater fentanyl intake than females (**Supplementary Figure 5**). This negative finding suggests that the decrease in fentanyl self-administration observed in sucrose-trained mice (**Supplementary Figure 5A-D**) could have been influenced by prior sucrose experience.

To further address fentanyl self-administration in experimentally-naïve mice, we examined fentanyl self-administration across a range of lower drug doses (0.01-3 mg/L). Neither Sex nor Genotype influenced the doseresponse functions for active or inactive nose-poking (see Supplemental Results; **Supplementary Figure 6**). However, there was a sex-dependent effect of *Hn1+/−* on the allocation of responding in the fentanyl-reinforced hole (Genotype × Sex: F_1,135_ = 7.02, p = 0.01). This interaction was not reflected in females (**Fig.4A**; Genotype effect: p = 0.90); rather, *Hn1*+/− males showed greater fentanyl-appropriate responding than WT males (**Fig.4B;** Genotype effect: F_1,15_ = 11.38, **\***p = 0.004). While the allocation of responding in the active hole also decreased as a function of fentanyl dose (Dose effect: F_4,140_ = 7.41, p < 0.0001), neither Sex nor Genotype influenced the shape of this dose-response function (**Fig. 4A,B**; Dose interactions, p’s > 0.75). Overall, males consumed more fentanyl than females during dose-response testing (**Fig.4C,D**; Sex effect: F_1,35_ = 11.14, p = 0.002; Sex × Dose: F_4,140_=9.61, p < 0.0001). Moreover, there was a Genotype × Sex interaction with respect to the dose-response function of fentanyl intake (Genotype × Sex × Dose: F_4,140_ = 2.44, p = 0.049). Deconstructing this 3-way interaction along the Sex factor indicated no genotypic difference in fentanyl intake in females (**Fig.4C**; Genotype effect and interaction, p’s > 0.88). In contrast, the dose-response function was shifted downwards in *Hn1*+/− males, relative to their WT male counterparts (Genotype × Dose: F_4,60_ = 3.67, p = 0.01), with *Hn1*+/− mice showing less fentanyl intake at the 3 mg/L concentration (**Fig.4D:** t_15_ = 2.10, **\***p = 0.05) and a trend towards lower intake at 1.0 mg/L (t_15_ = 2.02, p = 0.06). Together, these latter findings indicate that *Hn1+/−* increases the reinforcing efficacy of fentanyl (as indicated by the shift upwards in fentanyl-appropriate responding), which likely reduces their propensity to consume fentanyl.

**Figure 4.**
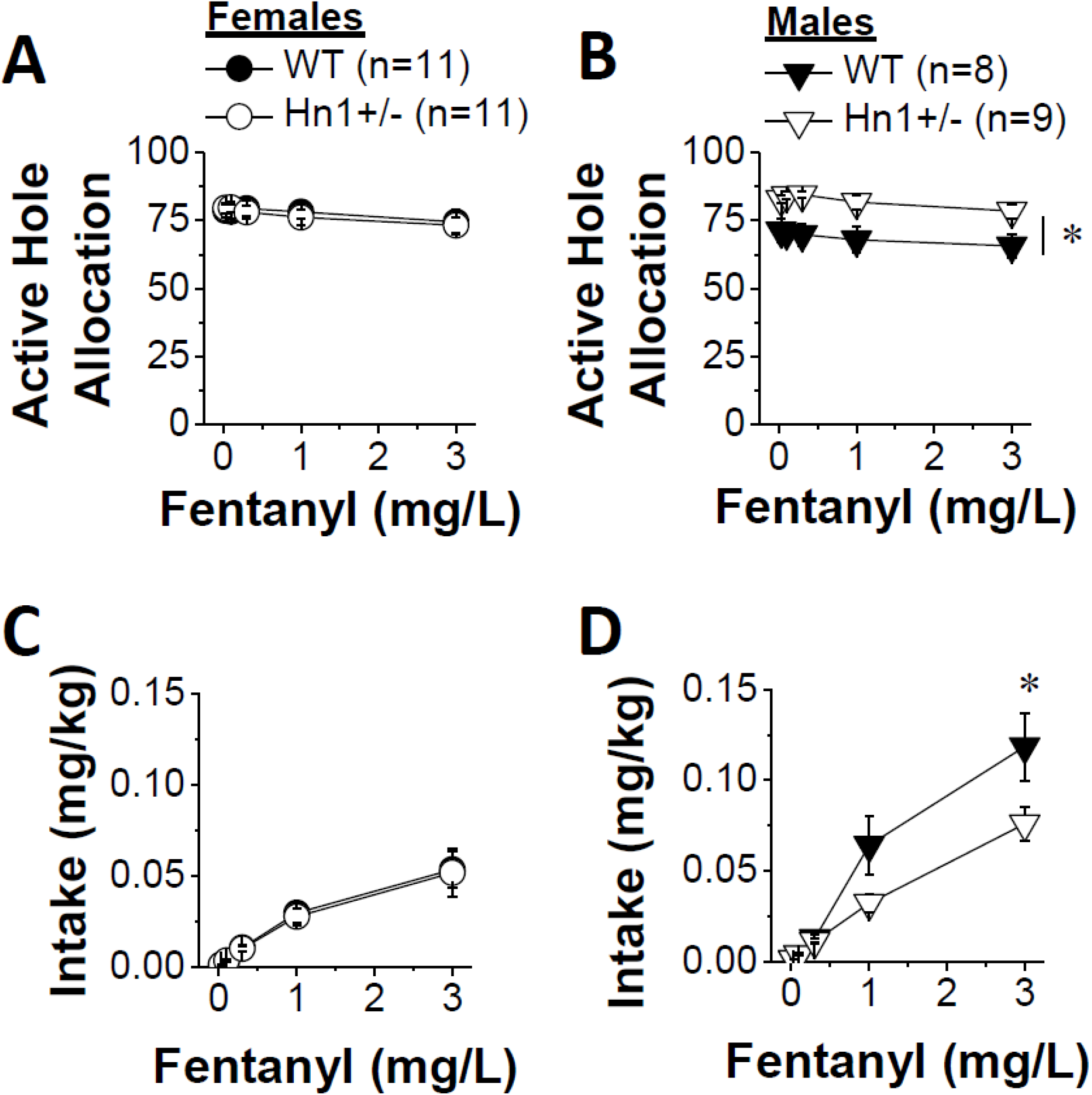
Sex-dependent effects of *Hn1*+/− on fentanyl reinforcement during acquisition of self-administration in sucrose-naïve mice. **(A):** When assessed across a range of low fentanyl doses (0.03-3.0 mg/L), no genotypic differences were apparent in experimentally-naïve females for the allocation of responding in the active, fentanyl-reinforced hole. (**B**): *Hn1*+/− males directed more nose-pokes toward the fentanyl-appropriate active lever, overall, than did their WT controls (**\***p = 0.004). **(C):** For females, there was no significant genotypic difference in low-dose FEN intake. **(D):** In contrast, *Hn1*+/− males consumed less FEN than WT males at 3 mg/L (**\***p = 0.05). Data represent the mean ± S.E.M. of the number of mice indicated in the figure legends. *p < 0.05 vs. WT; * denotes main effect of Genotype.

### Fentanyl withdrawal does not induce obvious sensorimotor-gating deficits in Hn1+/− or WT mice

As detailed in the **Supplementary Information**, neither *Hn1*+/− nor fentanyl withdrawal altered startle amplitude (**Supplementary Figure 7A**) or the pre-pulse inhibition (**PPI**) of startle amplitude (**Supplementary Figure 7B**). However, fentanyl-withdrawn males showed an overall modest impairment in PPI at both decibel levels (**Supplementary Figure 7C**).

### Sex-dependent modulation of behavior in light-dark shuttle box in Hn1+/− mice

Repeated fentanyl administration (twice daily injections of 0.8 mg/kg for 5 days) did not alter the number of light-side entries exhibited by either male or female WT or *Hn1*+/− mice (Drug effect and interactions, p’s > 0.13). However, a significant Genotype × Sex interaction was detected for this measure (F_1,100_ = 4.68, p = 0.03), that reflected a greater number of light-side entries in *Hn1*+/− males versus WT males (**Fig.5A:** t_49_ = 2.03, **\***p = 0.04), but no genotypic difference in female subjects (t-test, p = 0.29). None of the independent variables influenced the latency to enter, nor the time spent in, the light-side (data not shown; Genotype × Sex × Drug ANOVAs, all p’s > 0.14). These data from the light-dark shuttle box do not support an increase in anxiety-like behavior during early fentanyl withdrawal under this treatment regimen, but suggest that *Hn1*+/− reduces some signs of anxiety-like behavior in male mice only.

**Figure 5.**
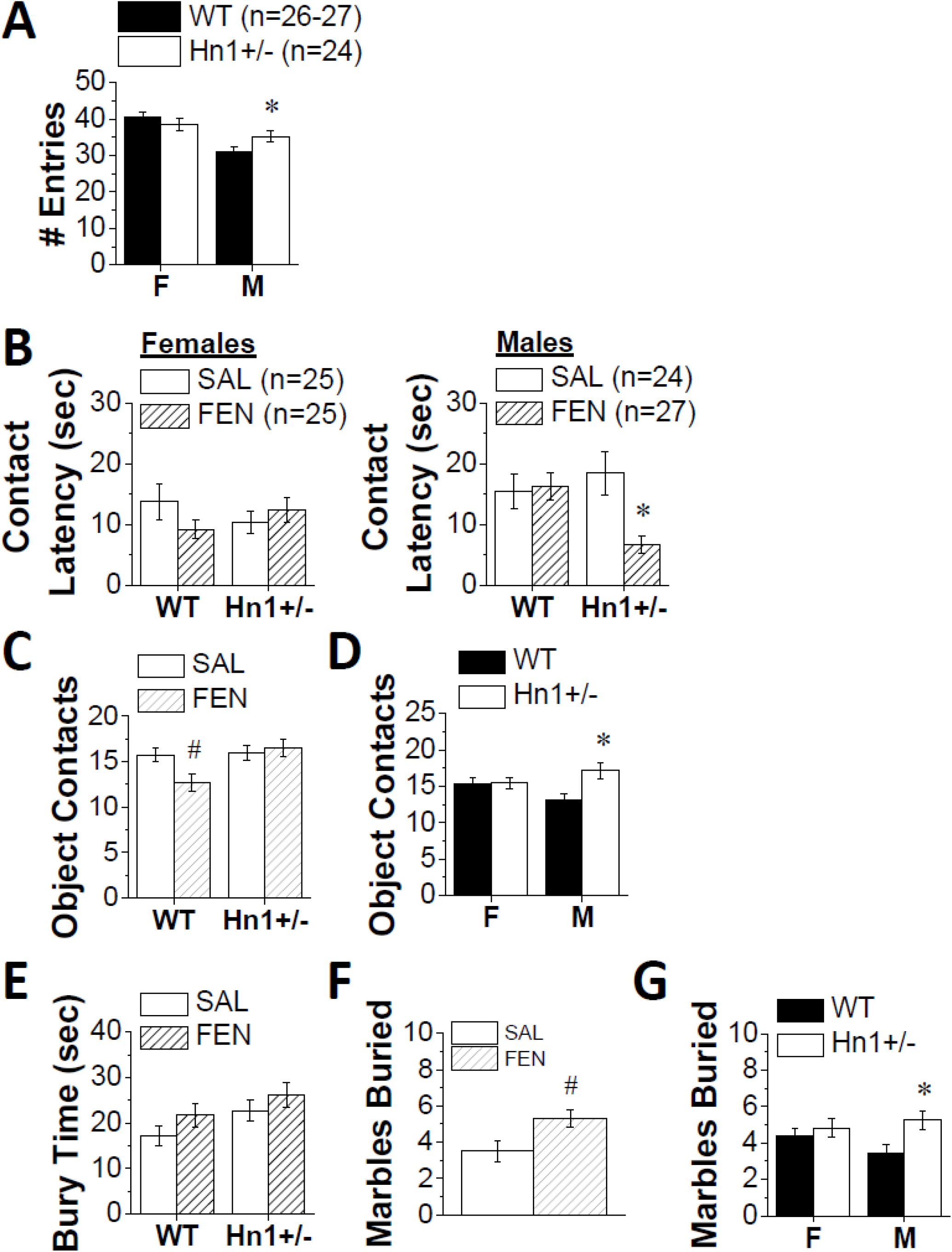
Sex-dependent effects of *Hn1*+/− on behavior in assays of anxiety-like behavior in naïve and fentanyl-withdrawn mice. **(A):** In the light-dark shuttle-box test, *Hn1*+/− males exhibited a greater number of light-side entries than WT males (**\***p = 0.04). **(B):** In the novel object encounter, fentanyl (**FEN**) withdrawal decreased the latency to first make contact with the object compared to saline (**SAL**) controls, but this effect was observed only in *Hn1*+/− males (right panel, versus WT males: **\***p = 0.002). **(C):** Irrespective of sex, fentanyl withdrawal reduced the number of novel object contacts in WT mice (**#**p = 0.01), but this effect was not apparent in *Hn1*+/− mice. **(D):** *Hn1*+/− males exhibited a higher number of contacts with the novel object than WT males (**\***p = 0.005), with no genotypic difference observed in females. **(E):** *Hn1*+/− mice trended towards spending more time burying marbles than WT mice and fentanyl withdrawal tended to increase the time spent burying. Neither of these main effects were statistically significant. **(F):** FEN withdrawal increased marble burying, irrespective of Genotype (**#**p = 0.01). **(G):** *Hn1+/−* males buried more marbles than WT males, irrespective of Treatment (**\***p = 0.01), with no genotypic difference observed in females. Data represent the mean ± S.E.M. of the number of mice indicated in the figure legends. **\***p < 0.05 vs. WT, **#**p < 0.05 vs. repeated SAL.

### Increased novel object approach behavior during fentanyl withdrawal in Hn1+/− males

Examination of the latency to first approach a novel object revealed a significant Genotype × Sex × Drug interaction (**Fig.5B**; F_1,100_ = 8.06, p = 0.006). Deconstruction of the interaction along the Sex factor failed to indicate any effect of Genotype or Drug on this measure in females (**Fig.5B**, left panel; Genotype × Drug interaction: p’s > 0.13). However, a significant Genotype × Drug interaction was detected in males (**Fig.5B**, right panel; F_1,150_ = 5.87, p = 0.02). As illustrated, this male-specific interaction reflected a shorter latency of fentanyl-treated *Hn1*+/− males to make contact with the novel object compared to WT males (**Fig.5B**, right; t_25_ = 3.51,**\***p = 0.002), while no genotypic difference was apparent for the contact latencies of saline control mice (t-test, p = 0.52). A significant Genotype × Drug interaction was also detected for the number of contacts with the novel object (F_1,100_ = 4.06, p = 0.05) that reflected a fentanyl-induced reduction in novel object contacts in WT mice (**Fig.5C**, left: t_48_ = 2.61, **#**p = 0.01), but no fentanyl effect in *Hn1*+/− mice (t-test, p = 0.70). There was also a significant Genotype × Sex interaction in the number of novel object contacts (F_1,100_ = 4.88, p = 0.03) that reflected a significant increase in *Hn1*+/− males relative to their WT male counterparts (**Fig.5D**; t_48_ = 3.00, **\***p = 0.005), but no genotypic difference in females (t-test, p = 0.94). None of our independent variables influenced the time in contact with the novel object (data not shown; Genotype × Sex × Drug ANOVAs, all p’s > 0.08). These data provide limited evidence that fentanyl withdrawal alters behavioral signs of negative affect in the novel object encounter assay. Further, these data provide additional evidence that *Hn1*+/− can increase approach behaviors in males, particularly those with a history of repeated fentanyl exposure.

### Fentanyl withdrawal increases marble burying behavior irrespective of genotype and Hn1+/− increases the number of marbles buried in males, irrespective of prior treatment

In examining marble-burying behavior during fentanyl withdrawal, there were no group differences in the latency to start marble-burying (data not shown; Genotype × Sex × Drug ANOVA, all p’s > 0.08). Overall, fentanyl withdrawal induced a modest, nonsignificant increase in time spent marble-burying (**Fig.5E**; Drug effect: F_1,100_ = 2.89, p = 0.09). While this fentanyl effect did not vary as a function of Genotype (Genotype × Drug: p = 0.71), *Hn1*+/− mice tended to spend more time burying marbles, overall, than their WT counterparts (**Fig.5E**; Genotype effect: F_1,100_ = 3.79, p = 0.06). Mice in fentanyl withdrawal buried a significantly greater number of marbles than saline-injected controls (**Fig.5F**; Drug effect: F_1,100_ = 6.61, p = 0.01), but this fentanyl effect did not vary as a function of either Sex or Genotype (Dose interactions, p’s > 0.14). However, there was a significant Genotype × Sex interaction observed for the number of marbles buried (F_1,100_ = 6.84, p = 0.01), which reflected a greater number of marbles buried by *Hn1*+/− males versus their male WT controls (**Fig.5G**; t_49_ = 2.66, p = 0.01), but no genotypic difference in females (**Fig.5G**; t-test, p = 0.41). Thus, as observed for alcohol withdrawal (e.g., Lee et al., 2017) the marble-burying test appears to be particularly sensitive at detecting an effect of early fentanyl withdrawal. However, in contrast to our other assays of anxiety-like behavior presented above, the data from this assay indicate an *anxiogenic-like* effect of *Hnrnph1* mutation that is selective for males.

### Fentanyl withdrawal induces a buspirone-reversible increase in swimming behavior in the forced swim test: No effect of Hn1 +/− on fentanyl withdrawal signs or the buspirone treatment response

In the forced swim test, fentanyl withdrawal increased the latency to first float, irrespective of the Genotype or Sex (**Fig.6A**; Drug effect: F_1,100_ = 6.87, **\***p = 0.01; all other p’s > 0.14). Fentanyl withdrawal also significantly reduced the time spent floating in a manner independent of Genotype or Sex (**Fig.6B**; Drug effect: F_1,100_ = 12.44, **\***p = 0.001; other p’s > 0.28), and a similar result was observed for the number of immobile episodes (**Fig.6C**; Drug effect: F_1,100_ = 16.74, **\***p < 0.0001; other p’s > 0.45]. These data indicate that, similar to alcohol withdrawal (Lee *et al.* 2015, 2016, 2017a), early withdrawal from repeated fentanyl administration induces a robust increase in swimming behavior in the forced swim test. However, *Hn1+/−* mutation does not alter this fentanyl withdrawal behavior in this assay.

**Figure 6.**
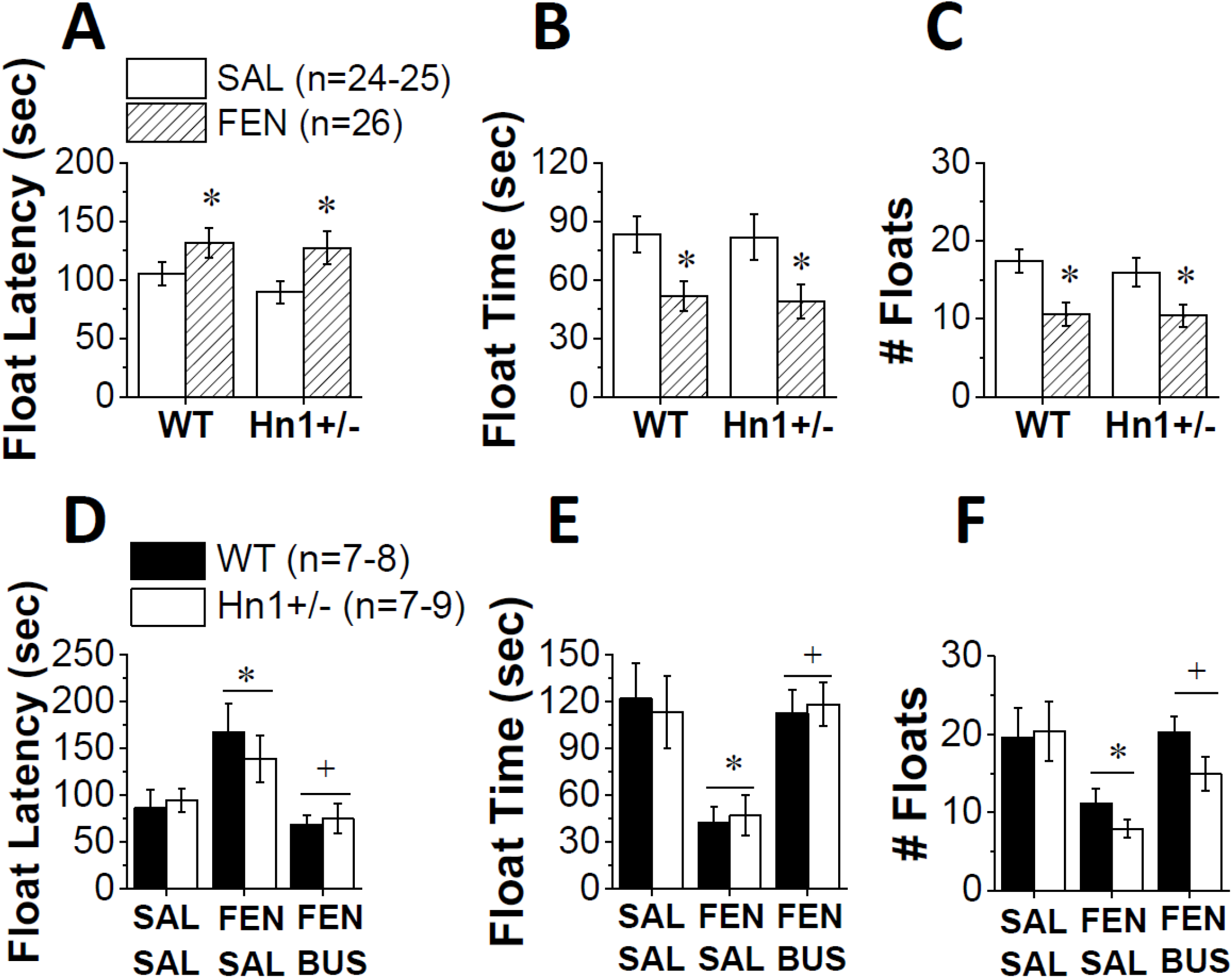
Buspirone pretreatment reverses the increase in swimming behavior in the forced swim test exhibited by WT and *Hn1*+/− mice during fentanyl withdrawal. Irrespective of genotype, fentanyl (**FEN**)- experienced mice exhibited **(A)** a longer latency to first float, **(B)** reduced time spent floating and **(C)** reduced number of floats, relative to their saline (SAL)-injected controls. In a separate cohort of mice, we replicated the effect of repeated fentanyl treatment upon swimming behavior (SAL-SAL vs. FEN-SAL). Importantly, pretreatment of a separate cohort of fentanyl-treated mice with 5 mg/kg buspirone (FEN-BUS) prior to testing reversed the fentanyl effect upon the latency to first float **(D)**, the time spent floating **(E)** and the number of floats **(F)**, when compared to their saline-pretreated controls (FEN-SAL). Data represent the mean ± S.E.M. **\***p < 0.05 vs. SAL or SAL-SAL; **+**p < 0.05 vs. FEN-SAL.

We next tested the hypothesis that the fentanyl withdrawal-induced increase in swimming observed in the forced swim test might reflect increased anxiety-like behavior by pretreating mice with an effective dose of the anxiolytic buspirone (Lee *et al.* 2017b). Replicating our above findings (**Fig.6A-C**), mice in early fentanyl withdrawal exhibited increased swimming, irrespective of Genotype or Sex (**\*Fig.6D-F**). As reported previously for alcohol withdrawal (Lee *et al.* 2017a), acute pretreatment with 5 mg/kg buspirone reversed the increased swimming behavior that was present in fentanyl-withdrawn mice (**Fig.6D-F**). This pattern of effects was statistically significant for the latency to first float (**Fig.6D**; Group effect: F_1,45_ = 7.57, **+**p = 0.002; LSD post-hoc tests: saline-saline vs. fentanyl-saline, p = 0.007; fentanyl-saline vs. fentanyl-buspirone, p = 0.001; no other main effects or interactions, p’s > 0.40), the time spent floating (**Fig.6E;** Group effect: F_1,45_ = 6.58, **+**p = 0.004; LSD post-hoc tests: saline-saline vs. fentanyl-saline, p = 0.001; fentanyl-saline vs. fentanyl-buspirone, p = 0.002; no other main effects or interactions, p’s > 0.34] and the number of floating episodes (**Fig.6F;** Group effect: F_1,45_ = 6.29, **+**p = 0.005; LSD post-hoc tests: saline-saline vs. fentanyl-saline, p = 0.001; fentanyl-saline vs. fentanyl-buspirone, p = 0.009; no other main effects or interactions, p’s > 0.25). These data replicate our original observation that opioid withdrawal-induced behaviors can manifest in the forced swim test as an increase in swimming and provide pharmacological validation that this swimming reflects anxiety-like behavior. However, *Hn1*+/− does not affect either the withdrawal or buspirone response.

### No changes in signs of fentanyl dependence in Hn1+/− mice

Although a modest sex difference was observed for some measures of physical dependence (**Supplementary Figure 8**), we detected no effects of *Hn1+/−* on naltrexone-precipitated withdrawal signs as detailed in the **Supplementary Information**.

## DISCUSSION

We previously linked *Hnrnph1 (**Hn1**)* polymorphisms to alterations in several methamphetamine-induced addiction-relevant behaviors in mice (Yazdani *et al.* 2015; Ruan *et al.* 2020a, 2020b). Furthermore, polymorphisms in *OPRM1* (mu opioid receptor) have been linked to changes in hnRNP H1 binding to and post-transcriptional processing of *OPRM1* (gene coding for mu opioid receptor) and heroin addiction severity in humans (Xu *et al.* 2014). Here, we demonstrate that one copy of a frameshift deletion in the first coding exon of *Hn1 (Hn1+/−)* can also alter opioid addiction-relevant behaviors in mice. Specifically, *Hn1*+/− mice showed reduced acute fentanyl-induced locomotor activity and low-dose fentanyl reward (CPP) and sex-interactive differences in fentanyl-induced locomotor sensitization (decreased in *Hn1*+/− females, increased in *Hn1*+/− males; **Figs. 1–2**). Furthermore, *Hn1*+/− mice showed reduced operant responding for sucrose (**Fig.3**) and subsequent reduced operant responding for fentanyl (**Supplementary Figure 4**). Sucrose-naïve *Hn1*+/− males showed evidence for increased efficacy of fentanyl reinforcement by demonstrating an increase in active hole allocation while at the same time showing reduced fentanyl intake (**Fig.4; Supplementary Figure 5**). Together, these results provide support that *Hn 1* dysfunction can alter behavioral responses to multiple classes of drugs of abuse.

The effects of *Hn1*+/− on fentanyl-induced behaviors were less pronounced compared to the effects on methamphetamine-induced behaviors (Yazdani *et al.* 2015; Ruan *et al.* 2020a). One reason could be related to the cellular mechanism proposed to underlie *Hn1*+/− modulation of drug-induced behaviors. *Hn1*+/− induced a profound reduction in methamphetamine-induced dopamine release in the nucleus accumbens as evidenced by *in vivo* microdialysis in the absence of any change in baseline extracellular dopamine levels, dopamine uptake, transporter expression, or transporter function (Ruan *et al.* 2020a). *Hn1*+/− also induced a two-fold increase in synaptosomal hnRNP H protein and proteomic analysis identified a highly enriched set of mitochondrial proteins that were perturbed at baseline and showed opposite methamphetamine-induced changes in synaptosomal expression/localization in *Hn1*+/− versus WT mice (Ruan *et al.* 2020a). Based on these observations, we proposed that increased synaptic hnRNP H in *Hn1*+/− mice alters mitochondrial gene expression and function to disrupt dopamine release and behavior in response to dopamine release-provoking stimuli. Methamphetamine as a stimulus acts directly at the site of action (dopamine transporters and vesicular monoamine transporters of the presynaptic dopaminergic neuron terminals) to cause a surge in extracellular dopamine levels. In contrast, like other opioids such as morphine (Johnson & North 1992), fentanyl is thought to disinhibit midbrain dopaminergic neurons to increase excitability, depolarization and dopamine release in the nucleus accumbens (Yoshida *et al.* 1999). Thus, if decreased dopamine release is the parsimonious mechanism that underlies reduced methamphetamine- and fentanyl-induced behaviors in *Hn1*+/− mice, a more pronounced behavioral effect would be expected in response to a psychostimulant compared to an opioid. It should also be noted that we previously found indirect evidence for modulation of hnRNP H staining in cultured rat primary neurons in response to D1 but not D2 receptor activation that was reversed by a D1 receptor antagonist (Ruan *et al.* 2018); thus, because our more recent study did not distinguish between the pre- and postsynaptic synaptosome (Ruan *et al.* 2020a), an alteration in post-synaptic D1 receptor signaling in *Hn1*+/− mice could also comprise a molecular mechanism underlying both psychostimulant and opioid behaviors (Awasaki *et al.* 1997).

What is the mechanism underlying the sex-interactive effects of *Hn1*+/− on behavior, in particular fentanyl-induced locomotor effects (**Fig.1**) and fentanyl reinforcement (**Fig.4; Supplementary Figure 5D**)? *Hnrnph2* is a gene homolog of *Hnrnph1* and is located on the X chromosome in both rodents and humans. Human mutations in *HNRNPH2* (and more recently *HNRNPH1)* have been linked to a rare, x-linked neurodevelopmental disorder in females (Bain *et al.* 2016; Pilch *et al.* 2018; Harmsen *et al.* 2019). Because males only have one copy of *Hnrnph2* whereas females have two, it is possible that *Hn1*+/− could lead to sex-dependent changes in *Hnrnph2* expression and behavior (in particular if *Hnrnph2* undergoes variable X-inactivation) that in turn, lead to sex-dependent differences in fentanyl-related behaviors. *Hn1*+/− is known to be associated with a dysregulation of dopaminergic function (Ruan *et al.* 2020a); thus, it is conceivable that a sex-specific change in *Hnrnph2* expression could alter the dopamine system and fentanyl-induced behaviors. Besides sex chromosome effects, a separate mechanism could involve sex steroids, which are known to modulate exogenous opioid-induced behaviors, including the discriminative stimulus and locomotor stimulant effects of morphine (Craft *et al.* 1999, 2006), as well as heroin and oxycodone self-administration (Becker & Chartoff 2019).

In agreement with our extensive previous behavioral battery in *Hn1*+/− mice, the behavioral effects of *Hn1*+/− in this study appear to be quite selective for drug-induced behaviors relevant to reward and reinforcement, with little evidence for effects on baseline behaviors. *Hn1*+/− mice showed unaltered nociception and fentanyl-induced antinociception after acute or chronic administration (**Fig.2; Supplementary Figure 3**). Furthermore, with the exception of a small but significant enhancement of marble burying behaviors in *Hn1*+/− males (**Fig.5**), *Hn1*+/− did not significantly alter several other behavioral measures, either in control mice or in mice receiving chronic fentanyl injections, including forced swim test behavior (**Fig.6**), startle (**Supplementary Figure 7**), or precipitated withdrawal (**Supplementary Figure 8**). These results provide further support that *Hn1*+/− exerts selective effects on drug-induced behaviors indicative of mesocorticolimbic dysfunction.

Another key finding of this study relates to the first report of an altered affective state during natural withdrawal from a repeated fentanyl injection regimen in mice on a C57BL/6J background. In humans, opioid withdrawal is associated with a number of affective or subjective signs, including dysphoria, irritability, and anxiety (Evans & Cahill 2016). For several decades, an opioid withdrawal-induced negative affective state has been successfully recapitulated in rodent models of morphine dependence using a variety of different behavioral assays (Hand *et al.* 1988; Bhattacharya *et al.* 1995; Grasing *et al.* 1996; Schulteis *et al.* 1998). More recently, genetic variability in the manifestation of oxycodone and fentanyl withdrawal-induced negative affect was highlighted through studies of different 129 mouse substrains (Jimenez *et al.* 2017; Szumlinski *et al.* 2019). Consistent with prior results for both opioid (Grasing *et al.* 1996; Jimenez *et al.* 2017; Szumlinski *et al.* 2019) and alcohol (Lee *et al.* 2015, 2016) withdrawal, the manifestation of the fentanyl withdrawal-induced negative affective state herein is task-specific. When assessed at 8-12 h following the last injection, fentanyl pre-exposed C57BL/6J mice made fewer novel objects contacts (**Fig.5C**), buried more marbles (**Fig.5F**) and exhibited more swimming behavior in the forced swim test (**Fig.6**) than saline-treated animals. In contrast, no fentanyl effect was apparent for acoustic startle (**Supplementary Figure 7**) or behavior expressed during the light-dark shuttle-box test in this study or that previous for 129 substrains (Szumlinski *et al.* 2019). The congruent findings across assays of neophobia/agoraphobia (Szumlinski *et al.* 2019), coupled with the reversal of the heightened swim reactivity in the forced swim test by the anxiolytic buspirone (**Fig.6D-F**), argues that our repeated fentanyl injection regimen is sufficient to elicit an anxiety-like state during early withdrawal. Buspirone has a complex pharmacology in that it acts as a partial agonist for 5-HT1A receptors to modulate monoamine release, in addition to antagonizing D2, D3 and D4 dopamine receptor subtypes (McMillen *et al.* 1983; Skolnick *et al.* 1984; Bergman *et al.* 2013). Furthermore, its metabolite acts as an alpha-2 adrenergic receptor antagonist (Gobert *et al.* 1999). The ability of acute buspirone to reverse the increased swimming behavior observed in opioid-withdrawn animals aligns with our prior results for alcohol withdrawal using this assay (Lee *et al.* 2017b). Such parallels in results suggest that opioid and alcohol withdrawal engage a common neurobiological mechanism that promotes maladaptive coping strategies related to augmented monoamine neurotransmission, which should be pursued in future work.

There are several limitations to this study. First, with regard to statistical power, we powered most of the experiments based on the effect size of the sex-combined effect of *Hn1*+/− deletion on methamphetamine-induced locomotor activity (Cohen’s d = 0.9; minimum of n=16 per genotype to achieve 80% power, p < 0.05) (Yazdani *et al.* 2015). In hindsight, the fentanyl effects were not as robust but nevertheless, we observed several instances of Sex × Genotype and Sex × Genotype × Treatment interactions. The reliability of these observations will need to be tested in a replication study with a larger, independent cohort. A second limitation is that our chronic fentanyl regimen did not induce robust signs of opioid tolerance. The lack of robust tolerance could be due to the rapid onset/offset of fentanyl physiological effects that dictate the time course of physiological recovery from opioid exposure and thus, the optimal time point for measuring physiologically perturbed behaviors. The subtle tolerance is unlikely to be explained by insufficient fentanyl dosing as 0.8 mg/kg is considered a high dose that induces clear, behavioral signs of intoxication – even one-half of this fentanyl dose (0.4 mg/kg) induces severe respiratory depression in C57BL/6J mice (Fechtner *et al.* 2015). Behavioral signs of fentanyl dependence (e.g., changes in elevated plus maze behavior) have been observed following repeated fentanyl dosing with 0.3 mg/kg in C57BL/6J mice but with the exception that a longer treatment regimen of two to four weeks was used (Fujii *et al.* 2019). Thus, a longer treatment regimen with 0.8 mg/kg fentanyl could induce more reliable opioid tolerance. Future studies will employ a different fentanyl regimen (longer treatment, shorter interval from final exposure to tolerance assessment) and additional opioids (morphine, oxycodone) to test for potential effects of *Hn1*+/− on opioid tolerance.

To summarize, we found decreased fentanyl-induced locomotor activity and reward in *Hn1*+/− mice as well as opposite effects on sensitization in *Hn1*+/− females (decrease) and *Hn1*+/− males (increase), and a male-selective enhancement of fentanyl reinforcing efficacy. In the context of our prior studies, these observations support a role for *Hn1* function in the behavioral response to both psychostimulants and opioids. By extension, we suspect that the behavioral and neurochemical effects of drugs of abuse from other classes and stimuli that are capable of robustly stimulating dopamine release will also be affected by *Hn1* dysfunction.

## ACKNOWLEDGEMENTS

The authors wish to thank Dylan Lieberman, Sami Ferdousian, and Nick Stailey for their assistance with video-scoring of mouse behavior. This work was supported by R01DA039168 (C.D.B.).

## DATA AVAILABILTY STATEMENT

All data in raw and processed forms will be made available upon request.

## SUPPLEMENTARY INFORMATION

### METHODS

#### Fentanyl antinociceptive tolerance

Experimentally naïve mice received twice daily i.p. injections of either saline (i.p.) or 0.8 mg/kg fentanyl (i.p.) for 5 consecutive days. On day 6, mice were tested for baseline nociception as described above. Thirty min post-baseline assessment, mice were injected with a single, challenge dose of fentanyl (0.4 mg/kg i.p.) and were tested on the hot plate for antinociception every 10 min for 1 h. Given the influence of social drug state on opioid behaviors (Eitan *et al.* 2017), mice within the same home cage all received the same treatment. Using R (https://www.r-project.org/), we analyzed fentanyl tolerance via mixed design ANOVA (between-subjects: Fentanyl Treatment, Genotype, Sex; within-subjects: Time).

#### Acoustic Startle

Testing was conducted in sound-attenuated startle chambers (SRLAB, San Diego Instruments, San Diego, CA). Each chamber comprised a Plexiglas cylinder (3.8 cm diameter) mounted on a Plexiglas platform, with a high frequency loudspeaker (28 cm above the cylinder) producing all acoustic stimuli. The background noise of each chamber was 70 dB. A piezoelectric accelerometer, attached to the base, detected and transduced movements within the cylinder were detected and transduced by a piezoelectric accelerometer and then digitized and stored by a PC-type computer. As conducted previously (Szumlinski *et al.* 2005), six different trial types were presented: startle pulse (st110, 110 dB / 40 milliseconds), low prepulse stimulus given alone (st74, 74 dB / 20 milliseconds), high prepulse stimulus given alone (st90, 90 dB / 20 milliseconds), st74 or st90 given 100 milliseconds before the onset of the startle pulse (pp74 and pp90, respectively) and no acoustic stimulus (i.e. only background noise was presented; st0). St100, st0, pp74 and pp90 trials were applied 10 times, st74 and st90 trials were applied five times, and all trials were given in random order. The average inter-trial interval was 15 s (10–20 s) and the data for startle amplitude were averaged across stimulus trials for each mouse prior to statistical analyses. Elevated basal activity and increased startle amplitude were interpreted as reflecting anxiety-like behavior, while reduced pre-pulse inhibition of acoustic startle was interpreted as reflecting a sensorimotor gating deficit. This test was approximately 20 min in length and immediately following testing, mice were removed from the startle chambers, the chambers and cylinders were wiped down with 30% ethanol, and mice were transported within the lab to a distinct testing room housing the shuttle boxes. The data for the individual startle stimuli were analyzed using a Genotype × Sex × Drug × Stimulus or Prepulse ANOVA, with repeated measures of the Stimulus (6 levels) or Pre-pulse (2 levels) factors.

#### Fentanyl physical dependence

The day following testing for negative affective-like behavior during opioid withdrawal, mice were injected in the morning with 0.8 mg/kg fentanyl and then, 8 h later, injected with 10 mg/kg naloxone to precipitate withdrawal. Behavioral signs of physical withdrawal were monitored for 30 min post-naloxone injection and then, at 1 h post-naloxone, mice were assessed for fecal and urine output, piloerection and reactivity to handling using a checklist. The data for each measure were analyzed by Sex × Genotype ANOVAs, using SPSS v.21 software.

### RESULTS

**Supplementary Figure 1:**
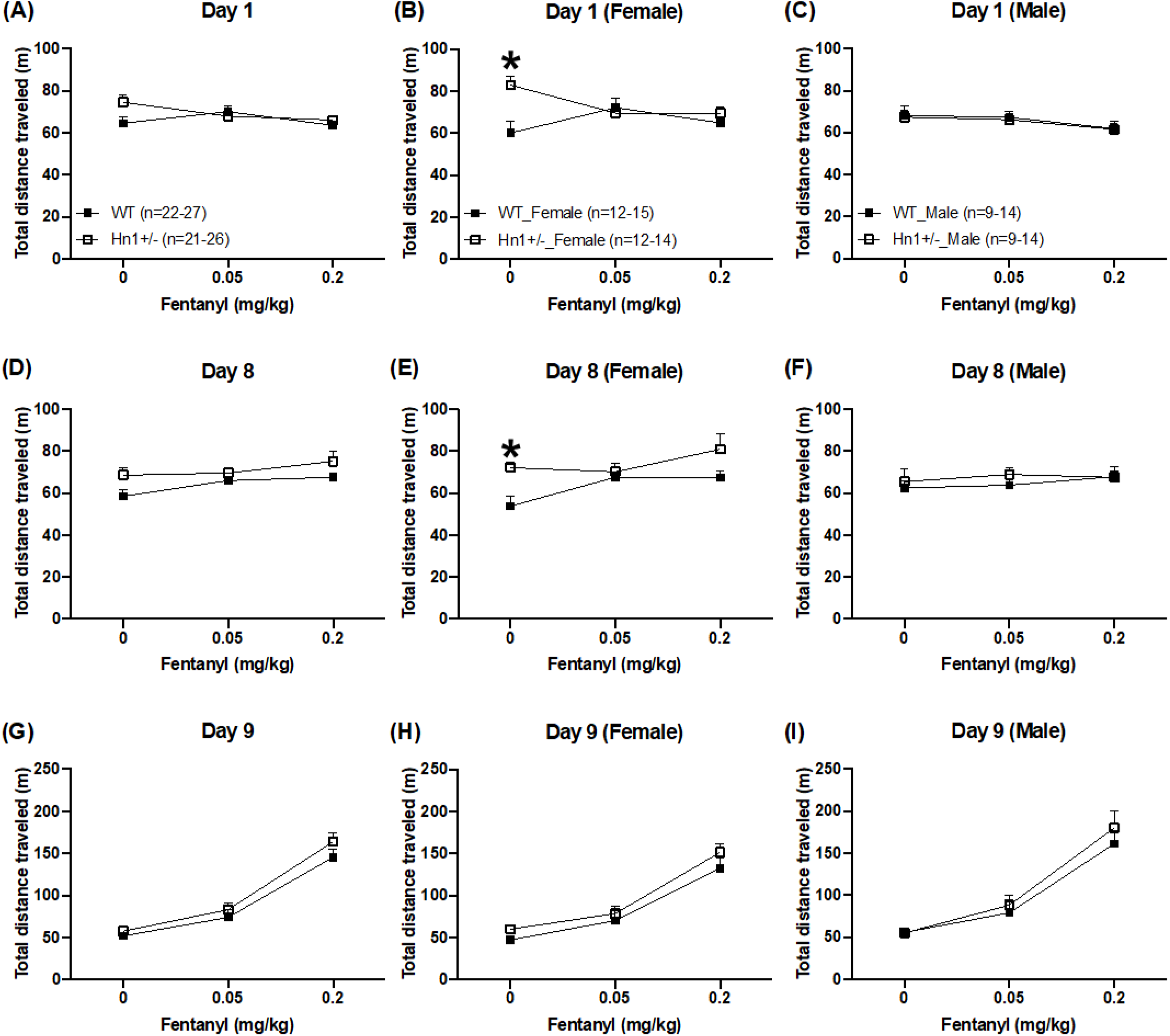
Locomotor activity during CPP assessment (Days 1, 8, and 9) **A-C:** For Day 1, there was a main effect of Sex (F_1,136_ = 4.06; p = 0.046), a Genotype × Sex interaction (F_1,136_ = 4.72; p = 0.032), and a Genotype × Sex × Dose interaction (F_2,136_ = 3.10; p = 0.048] that was explained by *Hn1*+/− females assigned the “0” dose (saline) showing (stochastically since treatment has not yet started) significantly greater locomotor activity than WT females (t_22_ = 3.36; *p =0.0028; **panel B**). **D-F:** For Day 8, there was a main effect of Genotype (F_1,136_ = 7.62; p = 0.0066) and Treatment (F_2,136_ = 3.33; p = 0.039) but no effect of Sex (F_1,136_ < 1) and no interactions (p’s > 0.08). The main effects of Genotype and Sex were driven by females assigned to the “0” mg/kg group (saline) whereby *Hn1*+/− females showed greater locomotor activity than WT females (t_22_ = 3.43; p = 0.0024; **panel E**). **G-I:** For Day 9, there was a main effect of Treatment (F_2,136_ = 96.09; p < 2.0 × 10^−16^) and Sex (F_1,136_ = 4.34; p = 0.039] but no effect of Genotype (F_1,136_ <1) or and no interactions (p’s > 0.19).

**Supplementary Figure 2:**
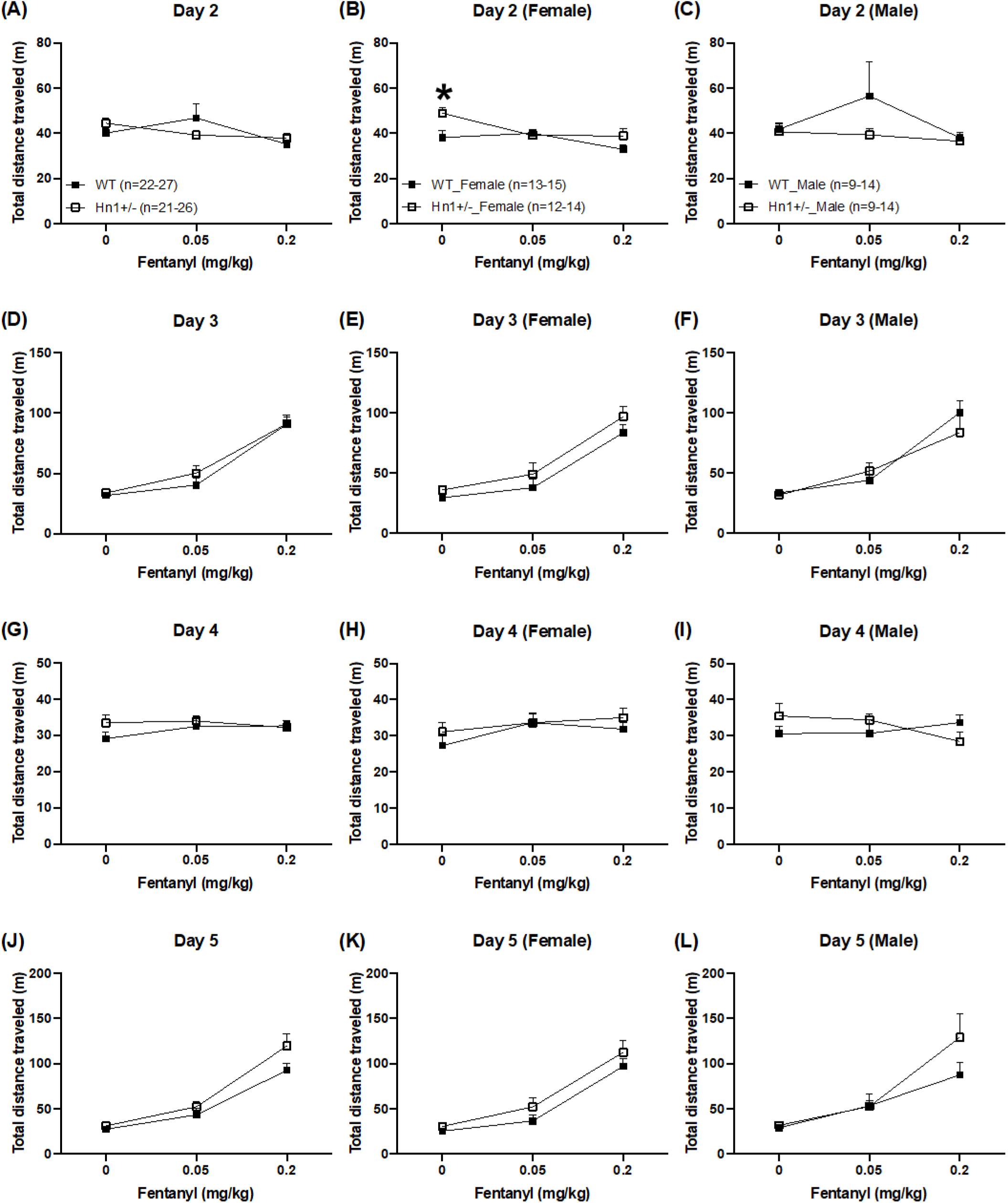
Locomotor activity during CPP training (Days 2, 3, 4, and 5) **A-C:** For Day 2, there was a (stochastic) main effect of Dose Assignment (F_2,136_ = 3.11; p = 0.048), a Genotype × Sex interaction (F_1,136_ = 6.08; p = 0.015), and a nonsignificant Genotype × Treatment interaction (F_2,136_ = 2.68; p = 0.073). There was no effect of Genotype or Sex (F_1,136_ < 1). Similar to Day 1, *Hn1*+/− females showed greater locomotor activity in the 0 mg/kg Treatment Assignment compared to WT females (t_22_ = 2.62; **\***p = 0.016; **panel B**). **D-F:** For Day 3, there was a main effect of Dose (F_2,136_ = 76.61; p < 0.0001) that was driven by the dosedependent increase in fentanyl-induced locomotor activity. There was no effect of Genotype (F_1,136_ < 1) or Sex (F_1,136_ < 1) and no interactions (p’s > 0.09). **G-I:** For Day 4, there was no effect of Genotype (F_1,136_ = 1.70; p = 0.2), Sex (F_1,136_ < 1), Treatment (F_2,136_ <1), or any interactions (p’s > 0.22). **J-L:** For Day 5, there was a main effect of Treatment (F_2,136_ = 67.92; p < 2.0 × 10^−16^) that was driven by the dose-dependent increase in fentanyl-induced locomotor activity (**panels J-L**) but no effect of Genotype (F_1,136_ = 2.35; p = 0.13), Sex (F_1,136_ < 1), or any interactions (p’s > 0.21).

**Supplementary Figure 3:**
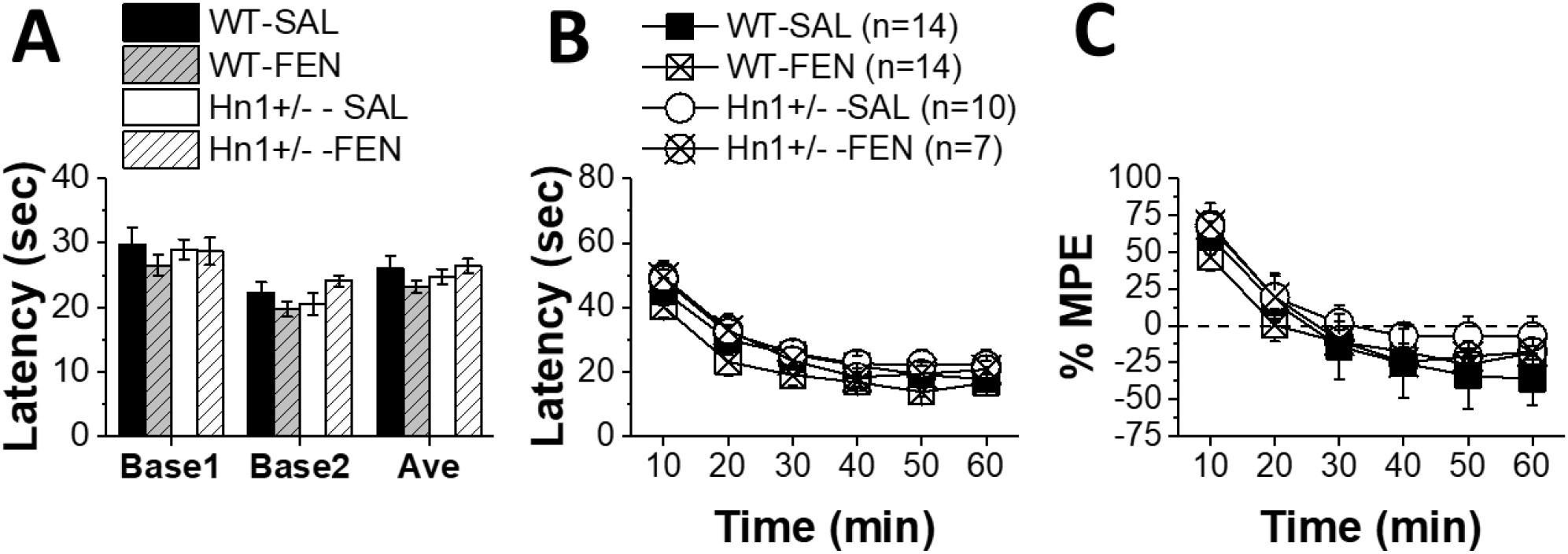
No difference in baseline nociception or in fentanyl-induced antinociception following chronic fentanyl administration in Hn1+/− mice. Following five days of twice daily injections of fentanyl (0.8 mg/kg, i.p.) or saline (i.p.), we assessed baseline nociception and the antinociceptive response to a fentanyl challenge injection (0.4 mg/kg, i.p.). **A:** For the first baseline nociceptive assessment, there was no main effect of Genotype (F_1,38_ < 1), Chronic Treatment (F_1,38_ = 1.07; p = 0.31), Sex (F_1,38_ < 1), or any interactions (ps > 0.07). For the second baseline nociceptive assessment, there was no effect of Genotype, Treatment, or Sex (F_1,38_ < 1), and no interactions (p’s > 0.05). For the average of the two baseline measurements, there was no effect of Genotype, Chronic Treatment, Sex (F_1,38_ < 1), or interactions (p’s > 0.18). **B:** In examining post-fentanyl latencies, there was no effect of Genotype (F_1,38_ = 2.33; p = 0.14), Treatment (F_1,38_ = 1.82; p = 0.19), Sex (F_1,38_ = 1.49; p = 0.23), and no interactions (p’s > 0.17). There was an effect of Time (F_5,190_ = 52.17; p < 0.0001) but no interactions with Time (p’s > 0.45). **C:** In examining % MPE, there was no effect of Genotype (F_1,38_ = 1.35; p = 0.25), Treatment, or Sex (F_1,38_ < 1) or any interactions (p’s > 0.27). There was an effect of Time (F_5,190_ = 33.07; p < 0.0001) but no interactions with Time (p’s > 0.39).

**Supplementary Figure 4:**
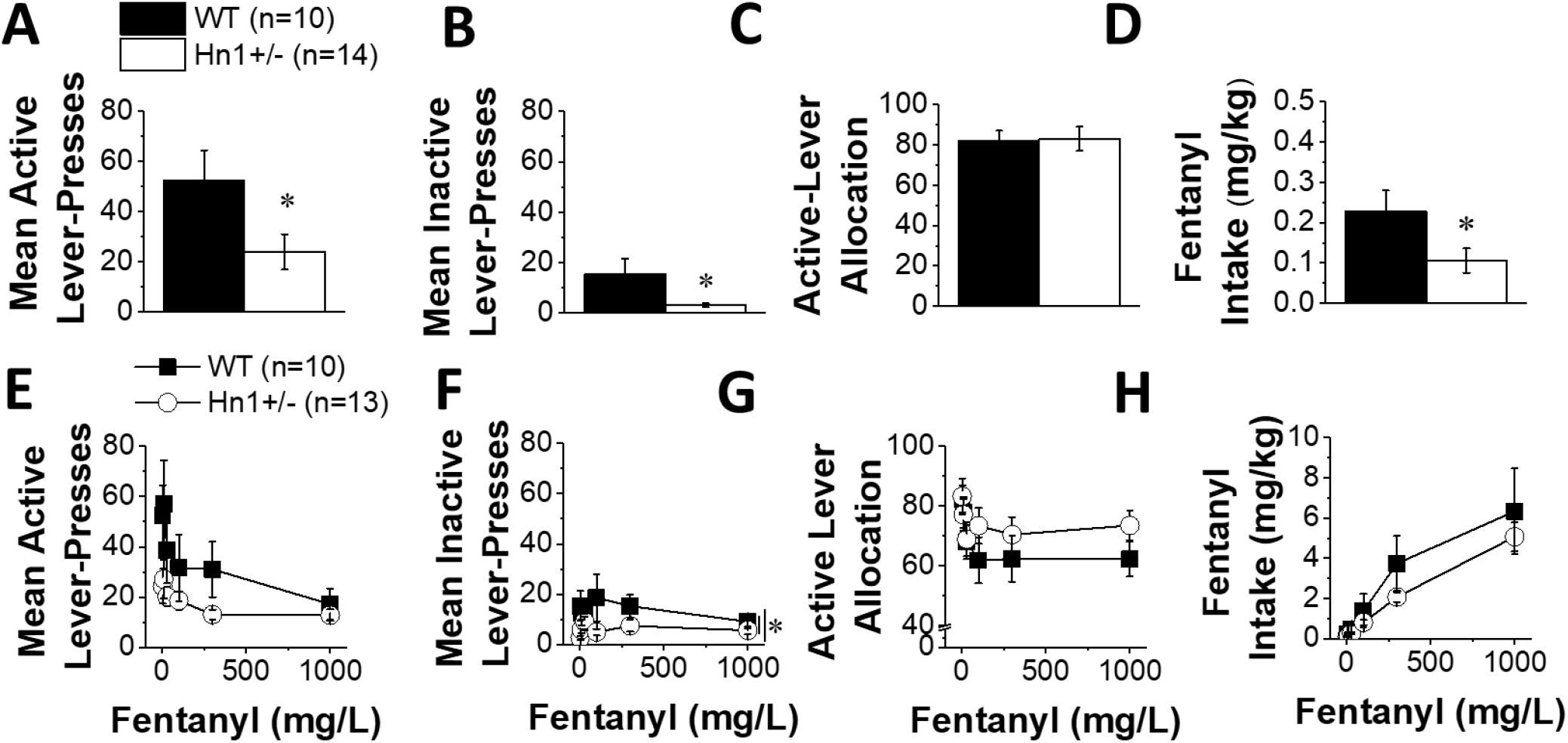
Deficits in fentanyl reinforcement/intake in Hn1+/− mice likely reflect a carryover effect from sucrose operant-conditioning. **(A):** When 3 mg/L fentanyl was substituted for sucrose, *Hn1*+/− mice perseverated in their lower level of active lever-responding (10-day period, t_22_ = 2.55, p = 0.02), although the magnitude of the genotypic difference in active lever-responding for 3 mg/L fentanyl was less robust at the end of this phase of training (last 3 days: t_22_ = 2.13, p = 0.04; WT = 52.27 ± 12.18 vs. *Hn1*+/− = 24.33 ± 6.89). **(B):** Consistent with their behavior during the acquisition of sucrose self-administration, *Hn1*+/− mice continued to exhibit a lower level inactive lever-responding upon 3 mg/L fentanyl substitution (for 10-day period, t_22_ = 3.44, p = 0.002; for last 3 days, t_22_ = 2.56, p = 0.02; WT = 15.43 ± 5.84 vs. *Hn1+/−* = 2.69 ± 0.84). **(C):** However, an analysis of their response allocation during this initial period of fentanyl self-administration failed to reveal genotypic differences in their response allocation for 3 mg/L fentanyl reinforcement (10-day period: t_22_ = 0.43, p = 0.67; last 3 days: t_22_ = 0.11, p = 0.91; WT = 82.32 ± 4.47% vs. *Hn1+/−* = 83.21 ± 5.99%). **(D):** Overall, *Hn1*+/− mice consumed less 3 mg/L fentanyl during initial training (10-day period: t_22_ = 2.16, p = 0.04); however, this genotypic difference in responding was no longer statistically significant by the end of the 3 mg/L fentanyl training period (t_22_ = 2.00, p = 0.06; WT: 0.23 ± 0.05 mg/kg vs. *Hn1*+/−: 0.12 ± 0.03 mg/kg). These data suggest that genotypic differences in fentanyl reinforcement/intake reflect carry-over effects from the sucrose-training phase of the experiment. **(E):** Consistent with this notion, *Hn1*+/− mice tended to lever-press less for fentanyl than WT mice during dose-response testing; however, the genotypic difference in active lever-responding for fentanyl was not statistically significant (Genotype effect: F_1,20_ = 3.82, p = 0.07; Genotype × Dose: F_5,100_ = 1.98, p = 0.09). Analysis of the dose-response function for active lever-presses revealed a descending curve (Dose effect: F_5,100_ = 8.73, p < 0.0001). **(F):** For the inactive lever-presses, the dose-response function was flat (**Fig. 4F**: Dose effect: F_5,100_ = 0.55, p = 0.74). Despite the lack of dose dependency for the inactive lever presses, *Hn1*+/− mice exhibited fewer inactive lever-presses than did WT mice (Genotype effect: F_1,20_ = 6.33, p = 0.02; interaction: F_5,100_ = 1.08, p = 0.38). **(G):** Examination of the dose-response allocation function revealed a clear dose-dependent reduction in the reinforcing properties of fentanyl (Dose effect: F_5,20_ = 3.75, p = 0.004); however, there was no evidence for any genotypic differences in this regard (Genotype effect: F_1,20_ = 0.48, p = 0.50; Genotype × Dose: F_2,100_ = 0.49, p = 0.79). **(H):** Analysis of the dose-intake function revealed a dose-dependent increase in daily fentanyl intake (Dose effect: F_5,100_ = 27.94, p < 0.0001) and although *Hn1*+/− mice tended to consume less fentanyl than their WT counterparts, no significant genotypic differences in fentanyl intake were detected (Genotype effect: F_1,20_ = 0.88, p = 0.36; interaction: F_5,100_ = 0.61, p = 0.69).

**Supplementary Figure 5:**
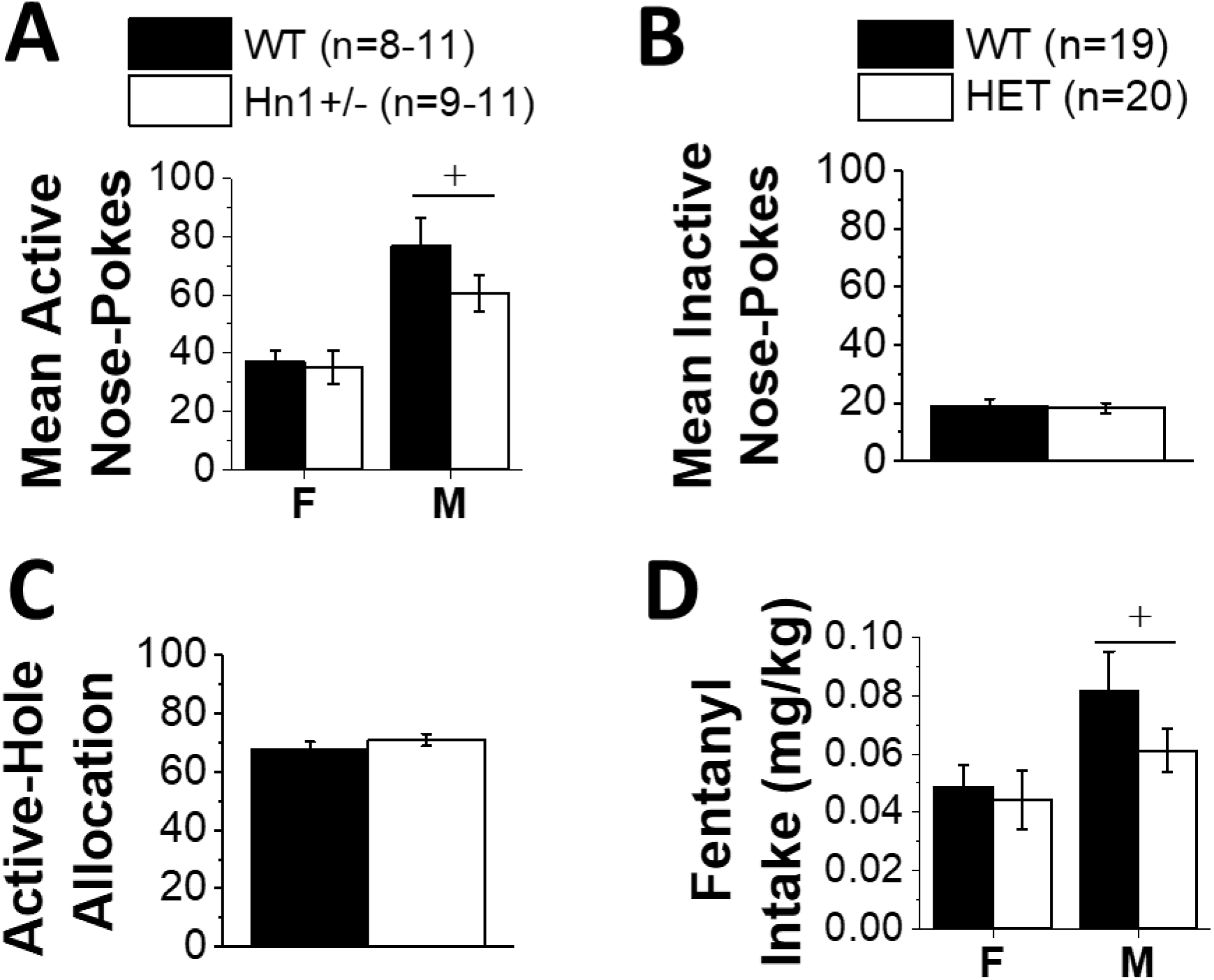
Fentanyl self-administration and intake at 3 mg/L in sucrose-naïve Hn1+/− and wild-type mice. In addition to testing the sucrose-experienced mice for fentanyl reinforcement and intake (**Supplementary Figure 2**), we trained a separate cohort of sucrose-naïve female and male mice to nose-poke for 3 mg/L fentanyl. **(A):** There was no genotypic difference in the average active responding over the 10 days of self-administration training, although males responded more than females, irrespective of Genotype (Sex effect: F_1,38_ = 15.05, **+**p < 0.0001; Genotype effect and interaction, p’s > 0.30). **(B):** Because there was no Sex or Genotype difference was noted for inactive hole pokes, we collapsed the data across Genotype for presentation (Genotype × Sex ANOVA, all p’s > 0.25). **(C):** Likewise, there were no sex or genotypic differences in the allocation of responding towards the active hole (Genotype × Sex ANOVA, all p’s > 0.08). **(D):** Notably, in the absence of sucrose-training, the average intake of the 3 mg/L fentanyl solution was approximately 25% of the average intake that was observed when mice were first trained to respond for sucrose reinforcement (compare **Supplementary Figure 3D versus Supplementary Figure 2D**). Males consumed more fentanyl than females during the training phase of this study (Sex effect: F_1,38_ = 6.53, **+**p = 0.02), but there was no genotypic difference in this regard (Genotype effect and interaction, p’s > 0.20). The lack of an effect of *Hn1+/−* on operant intake and consumption of 3 mg/L fentanyl in sucrose-naïve mice suggests that the moderate genotypic differences observed in the initial fentanyl self-administration study could reflect a carry-over effect from sucrose-training.

**Supplementary Figure 6:**
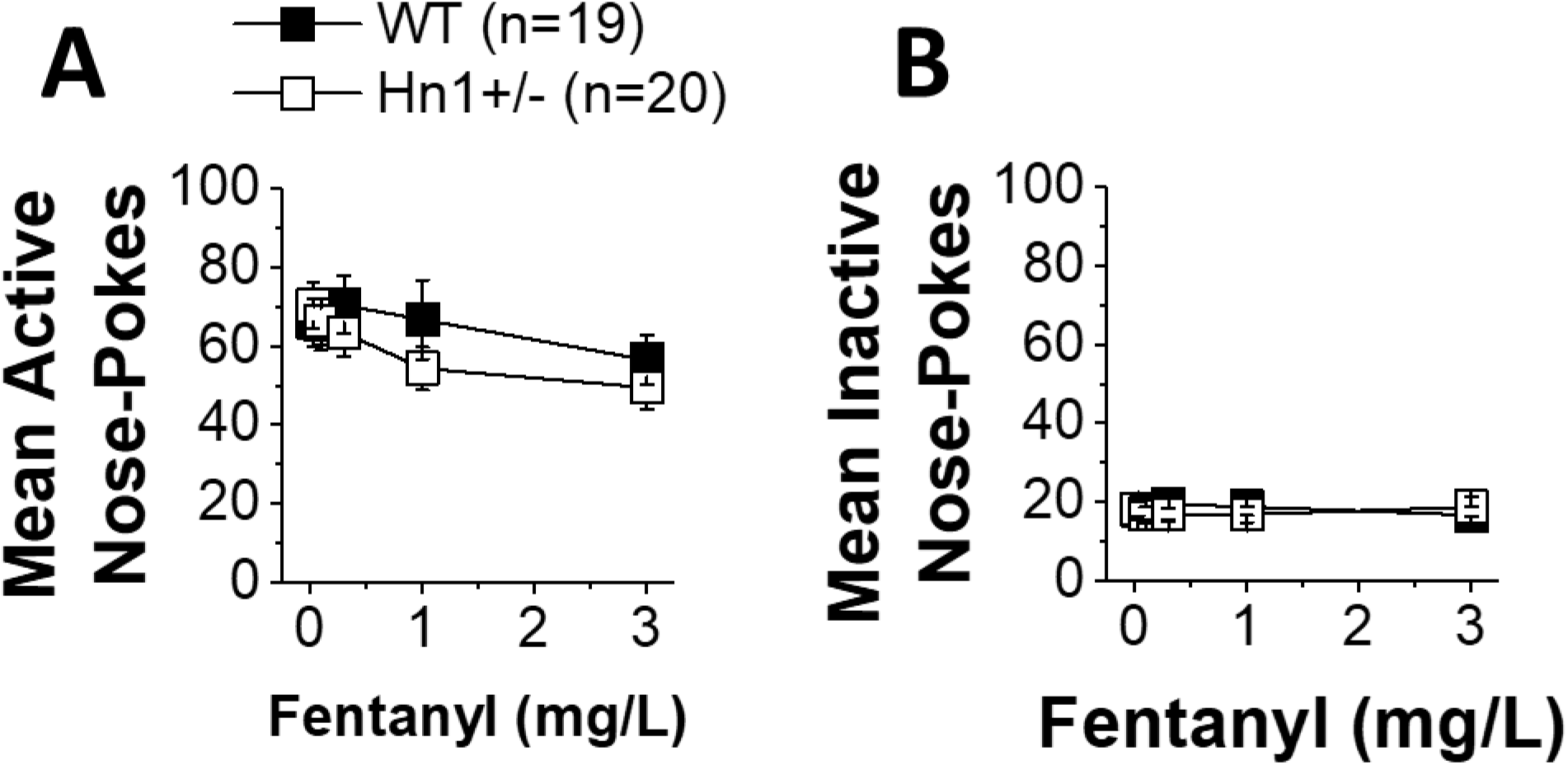
Active versus inactive nose pokes in a lower fentanyl dose range. **(A):** When tested across different low doses of fentanyl (0.03 – 3 mg/L), males continued to exhibit more nose-poking behavior than did females (data not shown) (Sex effect: F_1,35_ = 15.40, p < 0.0001; Sex × Dose: p = 1.00). Even when offered relatively low fentanyl doses, responding declined with increasing dose (Dose effect: F_4,140_ = 7.46, p < 0.0001) and a genotypic difference was detected by ANOVA (Genotype × Dose: F_4,140_ = 2.49, p = 0.04). However, post-hoc comparisons of the active hole-pokes did not indicate any significant genotypic differences at any of the fentanyl concentrations tested (t-tests, p’s > 0.25). **(B):** In contrast, nose-pokes in the inactive hole did not vary as a function of fentanyl dose, nor was this measure influenced by Sex or Genotype (Genotype × Sex × Dose ANOVA, all p’s > 0.20).

**Supplementary Figure 7:**
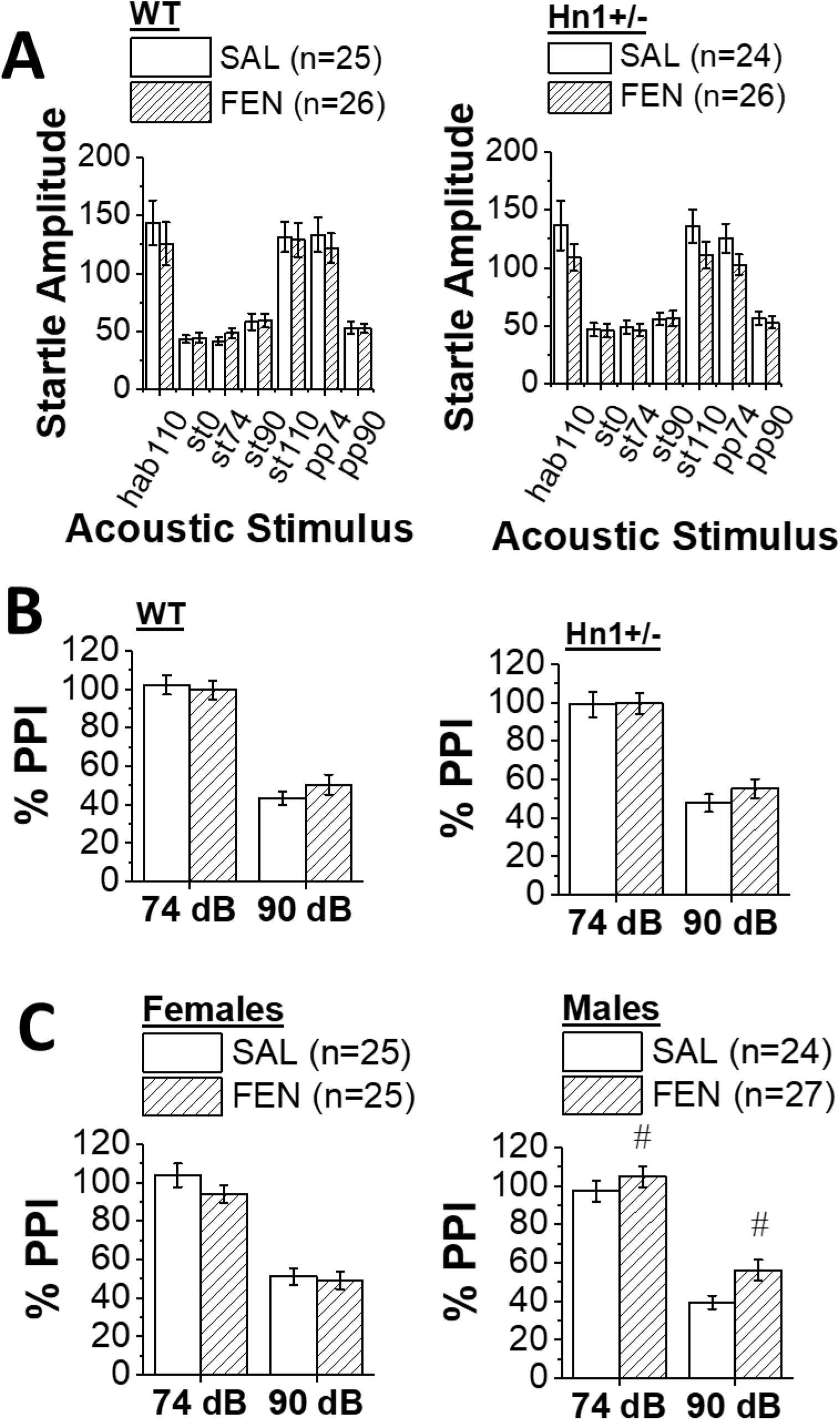
Hnrnph1 deletion does not impact acoustic startle or PPI. **(A):** Analysis of the startle amplitude elicited by each of the different acoustic stimuli indicated no effect of Genotype or interaction with Sex (p’s > 0.15). However, there was a significant Sex × Drug × Tone interaction (F_6,180_ = 3.64, p = 0.002). Deconstruction of this interaction along the Tone factor revealed sex differences in the effects of fentanyl for the 110 dB startle stimulus (Sex × Drug: F_1,37_ = 6.79, p = 0.01), which reflected a fentanyl-induced reduction in startle responsiveness in males (t_15_ = 2.14, p = 0.04), but not in female mice (t-test, p = 0.26). **(B):** The magnitude of PPI was not affected by early fentanyl withdrawal (open vs. closed; no Drug effect or interactions, p’s > 0.50), although a modest Sex × Genotype × Pre-pulse interaction was observed (F_1,30_ = 3.91, p = 0.06). Post-hoc analyses did not indicate any genotypic differences in males or females with respect to the PPI elicited by the 74 dB or 90 dB pre-pulses (left vs. right; t-tests, p’s > 0.20). **(C):** We also detected a significant Sex × Drug interaction with regard to PPI (F_1,30_ = 14.09, p = 0.001), which reflected a fentanyl-induced impairment in PPI magnitude in males (right panel) (for 74 dB, t_15_ = 2.22, **#**p = 0.04; for 90 dB, t_15_ = 3.23, **#**p = 0.006], but not in females (left panel; t-tests, p’s > 0.14). While these data indicate that males are more sensitive than females to fentanyl-induced disruptions in sensorimotor gating, they provide little support that disrupting *Hnrnph1* function perturbs basal or fentanyl-induced changes in acoustic startle or PPI.

**Supplementary Figure 8:**
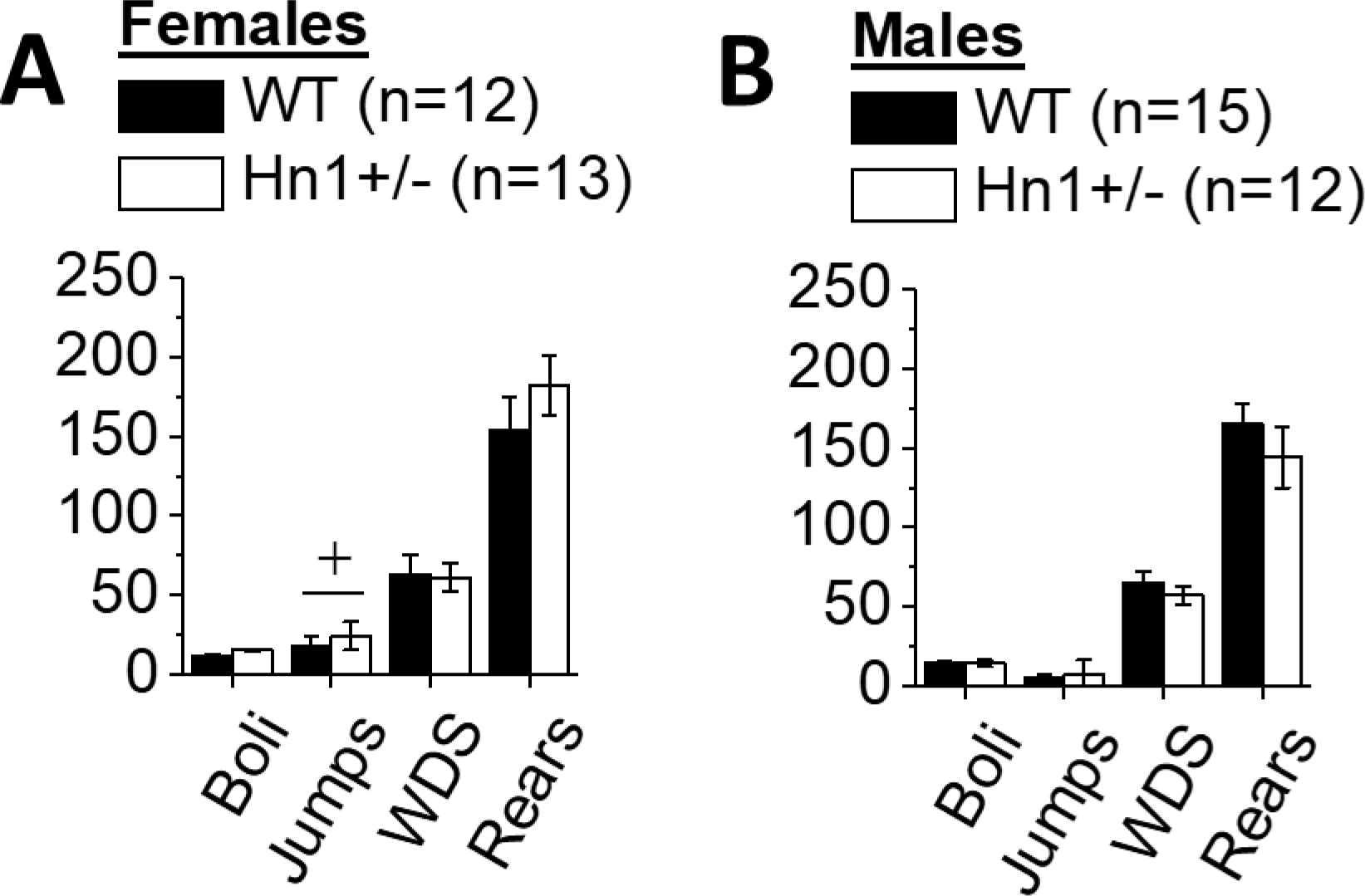
*Hnrnph1* deletion does not impact signs of fentanyl physical dependence. As expected, female mice weighed less than male mice at the outset of testing for precipitated withdrawal, irrespective of Genotype (Sex effect: F_1,51_ = 75.31, p < 0.0001; Genotype effect and interactions, p’s > 0.50). Following naloxone injection, females lost less weight than males by the end of the 1 h testing period, irrespective of genotype [Sex effect: F_1,51_ = 5.80, p = 0.02; no Genotype effect or interactions, p’s > 0.35; males: 0.99 ± 0.08 g (n = 27) vs. females: 0.67 ± 0.09 (n = 25)]. **(A,B):** In examining behavioral signs of opioid physical dependence, while females exhibited significantly more jumps than males (Sex effect: F_1,51_ = 6.81, **+**p = 0.01), there were no genotypic differences in this regard (Genotype effect and interaction, p’s > 0.45). None of the independent measures influenced fecal boli count (Genotype × Sex ANOVA, all p’s > 0.12), the number of wet dog shakes (all p’s > 0.55) or the number of rears (all p’s > 0.18) exhibited by the mice in response to naloxone. Further, no genotypic differences were observed for the presence of piloerection (λ^2^ = 0.07, p = 0.80), or diarrhea (λ^2^ = 0.58, p = 0.44) and all mice in the study urinated during testing. These data provide little evidence of fentanyl physical dependence under this regimen and at this time point post-chronic fentanyl administration. Furthermore, *Hn1+/−* does not alter behavioral or physiological signs of fentanyl dependence under these conditions.

